# Regeneration in the absence of a blastema requires cell division but is not tied to wound healing in the ctenophore *Mnemiopsis leidyi*

**DOI:** 10.1101/509331

**Authors:** Julia Ramon-Mateu, Tori Ellison, Thomas E. Angelini, Mark Q. Martindale

## Abstract

**Background:** The ability to regenerate is a widely distributed but highly variable trait among metazoans. A variety of modes of regeneration has been described for different organisms, however, many questions regarding the origin and evolution of these strategies remain unanswered. Most species of ctenophore (or “comb jellies”), a clade of marine animals that branch off at the base of the animal tree of life, possess an outstanding capacity to regenerate. However, the cellular and molecular mechanisms underlying this ability are unknown. We have used the ctenophore *Mnemiopsis leidyi* as a system to study wound healing and adult regeneration and provide some first-time insights of the cellular mechanisms involved in the regeneration of one of the most ancient extant group of multicellular animals.

**Results:** We show that cell proliferation is activated at the wound site and is indispensable for whole-body regeneration. Wound healing occurs normally in the absence of cell proliferation forming a scar-less wound epithelium. No blastema-like structure is generated at the cut site, rather undifferentiated cells assume the correct location of missing structures and differentiate in place. Pulse-chase experiments and surgical intervention show that cells originating in the main regions of cell proliferation (the tentacle bulbs) do not seem to contribute to the formation of new structures after surgical challenge, suggesting a local source of cells during regeneration. While exposure to cell-proliferation blocking treatment inhibits regeneration, the ability to regenerate is recovered when the treatment ends (days after the original cut), suggesting that ctenophore regenerative capabilities are constantly ready to be triggered and they are somehow separable of the wound healing process.

**Conclusions:** Ctenophore regeneration takes place through a process of cell proliferation-dependent non blastemal-like regeneration and is temporally separable of the wound healing process. The remarkable ability to replace missing tissue, the many favorable experimental features (e.g. optical clarity, high fecundity, rapid regenerative performance, stereotyped cell lineage, sequenced genome), and the early branching phylogenetic position in the animal tree, all point to the emergence of ctenophores as a new model system to study the evolution of animal regeneration.

## BACKGROUND

Regeneration, the ability to re-form a body part that has been lost, is a widely shared property of metazoans (1). However, the contribution of cell proliferation, the source of regenerating tissue, and the mechanisms which pattern the replaced tissues vary greatly among animals with regenerative ability, resulting in a collection of different “modes” of regeneration (2, 3). Based on the involvement of cell proliferation, there have been described cases of regeneration in which the restoration of the missing body part is accomplished in the absence of cell proliferation, by the remodeling of pre-existing cell tissues; on the other hand, and also more common, is the strategy of regeneration which requires active cell proliferation. Cell proliferation-based regeneration can involve the production of a regeneration-specific structure, the blastema. Several different definitions of a “blastema” have been proposed depending on the model system and its regenerative strategy. For example, in amphibians, the blastema is described as an unpigmented outgrowth consisting of a mass of undifferentiated progenitor cells that forms at the wound site from where cells proliferate and differentiate to form the missing structures (4, 5); while in planarians, the blastema is composed of post-mitotic progeny of proliferating cells that differentiate to reform the lost tissue (6). Given this lack of consensus around the word “blastema” and based on the biology, morphological features and regenerative response of our organism of study, we define the regeneration blastema as a “field” of undifferentiated cells that accumulate at the wound site and are later patterned to give rise to the appropriate set of missing structures but remains agnostic about their origin or their proliferative status.

The classical example of cell proliferation-independent regeneration is provided by the freshwater cnidarian polyp *Hydra*, which is able to regenerate the head after decapitation through remodeling of the pre-existing tissue without a significant contribution from cell proliferation (7–11). While documented cases of strict cell proliferation-independent regeneration are very few, most of the organisms with regenerative potential rely on cell proliferation – or a combination of both cell proliferation and tissue remodeling – to re-form lost structures. Regenerative abilities also appear to be diverse even within individual evolutionary clades. For example, regeneration of oral structures in another member of the phylum Cnidaria – *Nematostella vectensis* – is characterized by high levels of cell proliferation thus, differing from the cell proliferation-independent regeneration potential in *Hydra* (12). In planarians, whole-body regeneration is accomplished by the proliferation of pluripotent stem cells (neoblasts), the only cells in the adult with proliferative potential, which form a mass of undifferentiated cells known as the regenerating blastema (13–15). Annelid regeneration provides examples of both blastema-based regeneration and tissue-remodeling based regeneration (16, 17), showing diversity within the Lophotrochozoa. Moreover, evidence of cell migration has been documented during regeneration of several annelid species such as the freshwater annelid *Pristina leidyi* (18) and the marine annelid worm *Capitella teleta,* in which local (proliferating cells close to the wound site) and distant (stem cell migration) sources of cells contribute to the formation of the regenerating blastema (19). Evidence of cell migration during regeneration is also provided by the hydrozoan *Hydractinia echinata* in which stem cells (i-cells) from a remote area migrate to the wound site and contribute in the formation of the blastema (20). In vertebrates, regenerative potential is limited primarily to the structural or cellular level. Urodele amphibians are known for being the only vertebrate tetrapods that can regenerate amputated limbs as adults. Similar to the previous examples of cell proliferation-based regeneration, they require cell proliferation and the formation of a blastema. However, the urodele blastema is not generated from or composed of cells of a single type, but consists of a heterogeneous collection of lineage-restricted progenitors (21).

Among the animals with impressive whole-body regenerative capabilities are lobate ctenophores (comb jellies), fragile holopelagic marine carnivores that represent one of the oldest extant metazoan lineages. Ctenophora is latin for “comb bearer”, referring to eight longitudinally oriented rows of locomotory ctene (or comb) plates which they coordinately beat to propel through the water column. Ctenphores possess a highly unique body plan characterized by a biradial symmetry (with no planes of mirror symmetry) and two epithelial layers: the ectoderm and the endoderm, separated by a thick mesoglea mostly composed of extracellular matrix, but also containing several types of individual muscle and mesenchymal cells. The oral-aboral axis is their major body axis and it is characterized by the mouth at one (oral) pole and the apical sensory organ containing a statocyst at the opposite (aboral) pole. Most ctenophores possess a pair of muscular tentacles that bear specialized adhesive cells called colloblasts, used to capture prey (22) (Figure 1C). One of the best studied species of ctenophores is the lobate ctenophore *Mnemiopsis leidyi,* which is emerging as a new model system in evolutionary-developmental biology (23–28). *M. leidyi*’s life cycle is characterized by a rapid development including a highly stereotyped cleavage program and two adult stages: the juvenile tentaculate cydippid, distinguishable for having a pair of long branching tentacles (Figure 1A, 1B), and the larger lobate adult form which possess two oral feeding lobes. A particular feature of ctenophore embryogenesis is that they undergo mosaic development, meaning that embryos cannot compensate for cells/structures derived from cells killed or isolated during early development. If blastomeres are separated at the two-cell stage, each will generate a “half-animal,” possessing exactly half of the normal set of adult features (29, 30). This lack of ability to replace missing parts during embryogenesis contrasts with the outstanding capacity to regenerate as adults. Both the tentaculate larval and lobate adult life stages of *M. leidyi* readily regenerate and are capable of whole-body regeneration from only a single body quadrant (30).

**Figure 1.**
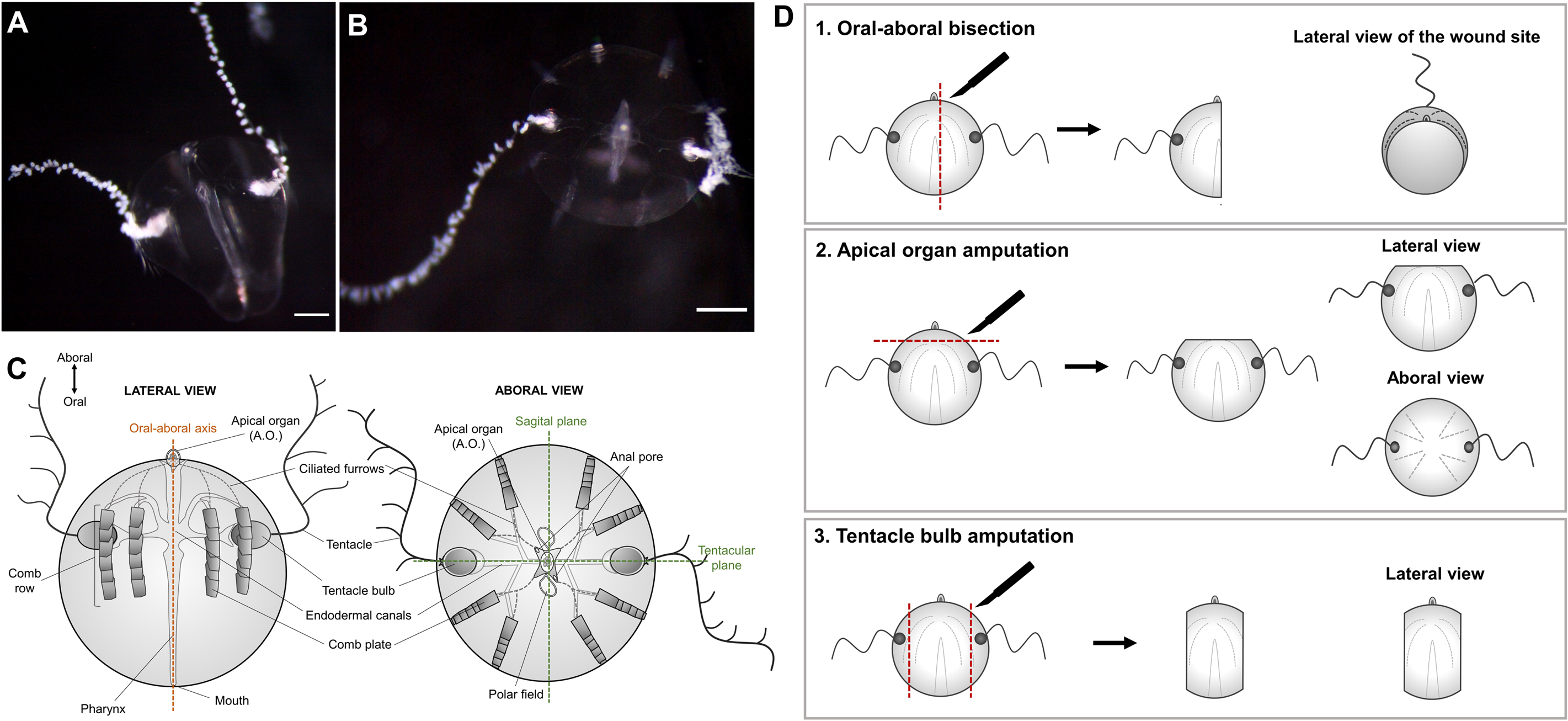
Cydippid stage of *Mnemiopsis leidyi* and animal surgeries. (**A**) Lateral view of a *M. leidyi* cydippid. (**B**) Aboral view of a *M.leidyi* cydippid. Scale bars = 100μm. (**C**) Schematic representation of the body plan of a cydippid stage in a lateral and aboral views. (**D**) Diagrams showing the three types of animal surgeries performed in this study and the views presented for each one.

It has been known for well over 80 years that ctenophores have the capacity to replace missing body parts (30–33) but the cellular and molecular mechanisms underlying this ability are poorly understood. Is cell proliferation required for ctenophore regeneration? Is any kind of blastema-like structure formed during regeneration? What is the source and nature of cells that contribute to the regenerated structures? What is the role of the wound epidermis in regulating the future regenerative outcome? We have studied wound healing and adult regeneration in the ctenophore *Mnemiopsis leidyi* and show that cell proliferation is activated at the wound site several hours after wound healing is complete and is indispensable for the regeneration of all the structures of the cydippid’s body. Wound healing occurs normally in the absence of cell proliferation forming a scar-less wound epithelium only a few hours after amputation. In both animals cut in half along the oral-aboral axis and those in which the apical organ is removed, anlage of all missing structures occurs within 24-48 hours and complete replacement of all cell types by 72 hours after the injury. No blastema-like structure is generated, rather undifferentiated cells assume the correct location of missing structures and differentiate in place. EdU (5-ethynyl-2’-deoxyuridine) labeling shows that in uncut animals the majority of cell divisions occur in the tentacle bulbs where the tentacles are continuously growing. In surgically challenged animals, cell division is stimulated at the wound site between 6-12 hours after injury and continues until 72 hours after injury. EdU pulse and chase experiments after surgery together with the removal of the two main regions of active cell proliferation suggest a local source of cells in the formation of missing structures. Although the appearance of new structures is completely dependent on cell division, surprisingly, the ability to regenerate is recovered when exposure to cell-proliferation blocking treatment ends, suggesting that the onset of regeneration is constantly ready to be triggered and it is somehow temporally separable from the wound healing process. This study provides some first-time insights of the cellular mechanisms involved in ctenophore regeneration and paves the way for future molecular studies that will contribute to the understanding of the evolution of the regenerative ability throughout the animal kingdom.

## RESULTS

### Whole-body regeneration in *Mnemiopsis leidyi* cydippids

Although the regenerative response has been studied previously in *M. leidyi* (23, 30–33) we first characterized the sequence of morphogenic events during cydippid wound healing and regeneration to provide a baseline for further experimental investigations. For this, two types of surgeries – representing the replacement of all the structures and cell types of the cydippid’s body (e.g. apical organ, comb rows, tentacle bulbs and tentacles) – were performed: 1) Bisection through the oral aboral axis keeping the piece with an intact apical organ which required the remaining piece to regenerate 4 comb rows and a tentacle apparatus (bisected cydippids with a complete intact apical organ regenerate into whole animals in a higher percentage of the cases compared to bisected animals with a half apical organ (30), 2) Apical organ amputation consisting in the removal of the apical organ, requiring the remaining piece to regenerate dome cilia, balancing cilia, lithocytes, polar fields, and neural cells of the apical organ floor (Figure 1D). The timing and order of formation of missing structures was assessed by *in vivo* imaging of the regenerating animals at different time points along the regeneration process.

#### Wound healing

To assess the mechanism of wound healing, juvenile cydippids were punctured generating a small epithelial gap (Figure 2A). (Imaging of larger wound healing events provided to be too difficult to document visually). Within minutes after puncture, the edges of the cells lining the gap increased their thickness indicating the start of the wound closure. The next phase of wound closure was characterized by the migration of a small number of cells coming from deep levels of the mesoglea (underneath the epithelial layer) to the edges of the wound (Figure 2B**, Additional file 1**). Interestingly, while the migration of cells from the mesoglea to the wound site was quite evident, the migration of epithelial cells across the wounded area was not observed. Once the migrating deep cells adhered to the edges of the gap, they started to extend spike-shaped cytoskeleton extensions (filopodia) laterally towards the adjacent cells. The diameter of the gap was progressively reduced as the connections between filopodia of marginal cells pulled the edges of the wound together (Figure 2C). When the diameter of the gap was significantly reduced, the cells at the margind of the gap started to extend filopodia not only to adjacent cells, but also to cells from the opposite edge of the wound. At this stage, multiple filopodia were detected emerging from a single cell (Figure 2D). Filopodia from all the edges of the wound eventually met forming a network of filaments that sealed the gap (Figure 2E) resulting in a scar-free epithelium within approximately 1 −1.5 hours after the procedure.

**Figure 2.**
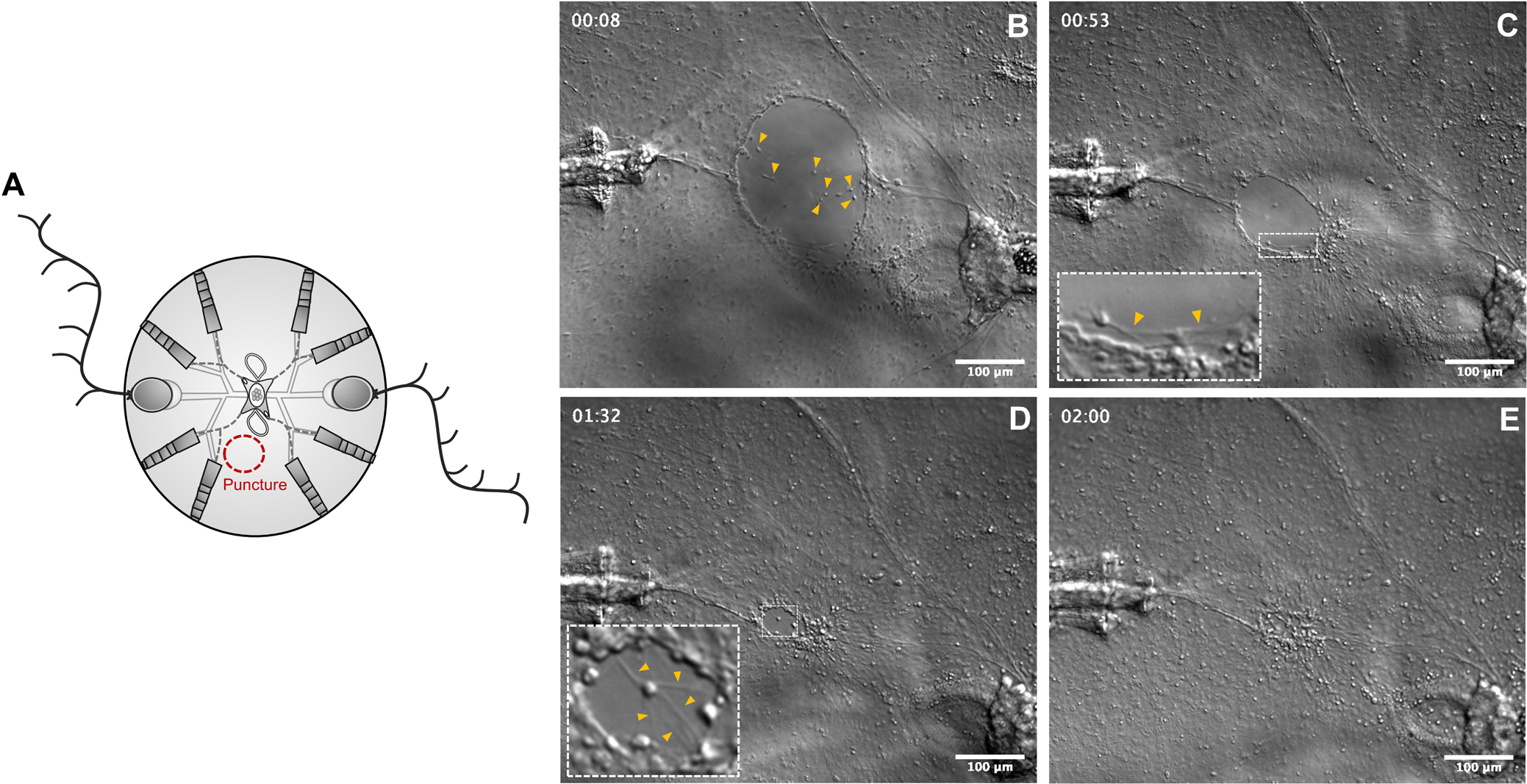
Wound healing by filopodia-dependent cell crawling. (**A**) Schematic representation of the puncture assay. (**B-E**) DIC images of the main phases of wound closure (See **Additional file 1** for the time lapse video corresponding to these images). (**B**) Cells from the mesoglea (yellow arrow caps) migrate upwards and adhere to the edges of the wound. (**C**) Marginal cells of the wound gap extend filopodia to the adjacent cells pulling the edges of the wound together. The inset shows a closer look to the cells at the edge of the wound and yellow arrow caps point to the filopodia. (**D**) When the diameter of the gap is considerably reduced, cells of the wound edge extend filopodia towards the opposite edges of the gap. The inset shows a cell extending multiple filopodia. (**E**) Network of filopodia connecting all the edges of the wound. Timescale is in hours. Scale bars = 100μm.

#### Events during half-body regeneration of Mnemiopsis leidyi cydippids following bisection through the oral-aboral axis

Cydippid stage animals were bisected through the oral-aboral axis with one half retaining the intact whole apical organ. Bisected cydippids containing half of the set of structures present in intact cydippids (four comb rows and one tentacle) and a complete apical organ were left to regenerate in 1x filtered sea water (1x FSW) (n>100) at 22°C. Wound closure was initiated rapidly after bisection with the edges of the wound forming a round circumference that continued to reduce in diameter until meeting and was completed within 2 hours after bisection (hab). No scar or trace of the original wound was evident after this time. About 16 hab, four ciliated furrows – which connect the apical organ with the comb rows – appeared on a surface epithelium at the aboral end of the cut site (Figure 3B). A large mass of cells (blastema-like structure) never appeared at the cut site. Rather, accumulations of cells were detected forming the primordia of all four of the future comb rows in a deeper plane at the end of each ciliated furrow. By 24 hab, the first comb plates appeared, first in the two most external (closer to existing comb rows) comb rows and later in the two internal rows (Figure 3C). Comb plate formation did not follow a consistent pattern initially. The correct orientation of comb plates and coordination of their beating was accomplished after a number of comb plates were formed (Figure 3D surface, deep), as has been described previously (33). Within 40 hab, coordinated comb plates were beating in all four regenerating comb rows and the primordia of tentacle bulb had emerged in the middle of the four comb rows. By 48 hab, regeneration of the missing structures of the cydippid body was essentially completed including the formation of the tentacle growing from the tentacle bulb (Figure 3E). At 96 hours after bisection, the regenerated tentacle was long enough to actively catch prey. The cut side continued to grow and within a day or two it was indistinguishable from the uncut side (Figure 3F).

**Figure 3.**
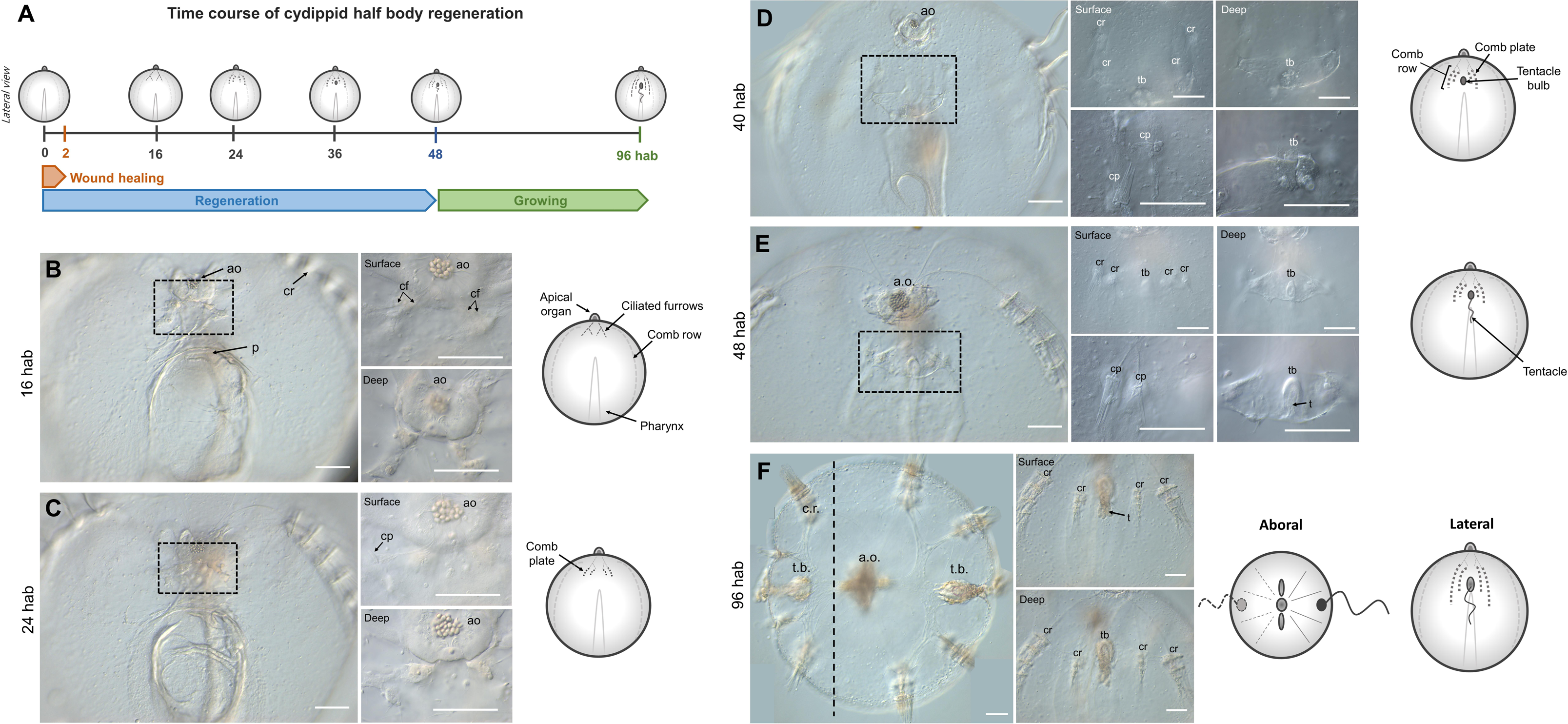
Half body regeneration in *Mnemiopsis leidyi* cydippid. (**A**) Schematic representation of the time course of morphogenic events during cydippid half body regeneration. All cartoons correspond to lateral views of the cydippid’s body bisected through the oral-aboral axis showing the cut site in the first plane. The apical organ is located at the aboral end (top) and the mouth at the oral end (down). For simplicity, tissues on the opposite body site are not depicted. (**B-F**) DIC images showing the cut site of regenerating cydippids from 16 to 96 hab. Dotted line rectangles in (**B**), (**C**), (**D**) and (**E**) show the area corresponding to higher magnifications on the right. Magnifications show surface and deep planes. The vertical dotted line in (**F**) indicates the approximate position of bisection, and all tissue in the left of the line is regenerated tissue. N>100. Scale bars = 100 μm. Abbreviations: hours after bisection (hab), apical organ (ao), pharynx (p), comb row (cr), ciliated furrow (cf), comb plate (cp), tentacle bulb (tb), tentacle (t).

#### Events during regeneration of Mnemiopsis leidyi cydippids following apical organ amputation

Cydippids in which the apical organ was amputated were left to regenerate in 1x FSW (n>100) at 22°C. The cut edges of the wound met and sealed within 30-60 minutes of the operation and the lesion was completely healed around 2 hours post amputation (hpa). Between 6 and 12 hpa, cells congregated under the wounded epithelium forming the primordia of the future apical organ (Figure 4E-F’). Extension of the ciliated furrows from each comb row towards the wound site could be spotted around 12 hpa. Within 24 hpa, cells at the wound site started to differentiate into the floor of the apical organ and its supporting cilia (Figure 4G-H’). At 48 hpa all the components of the statolith, including the supporting cilia, the balancing cilia and lithocytes, were formed (Figure 4I-J’). At approximately 60 hpa the complete set of structures forming the apical organ were regenerated with the exception of the polar fields (Figure 4K-L’). Within 3 days after surgery, the polar fields had formed, and animals were indistinguishable from control animals of the same size.

**Figure 4.**
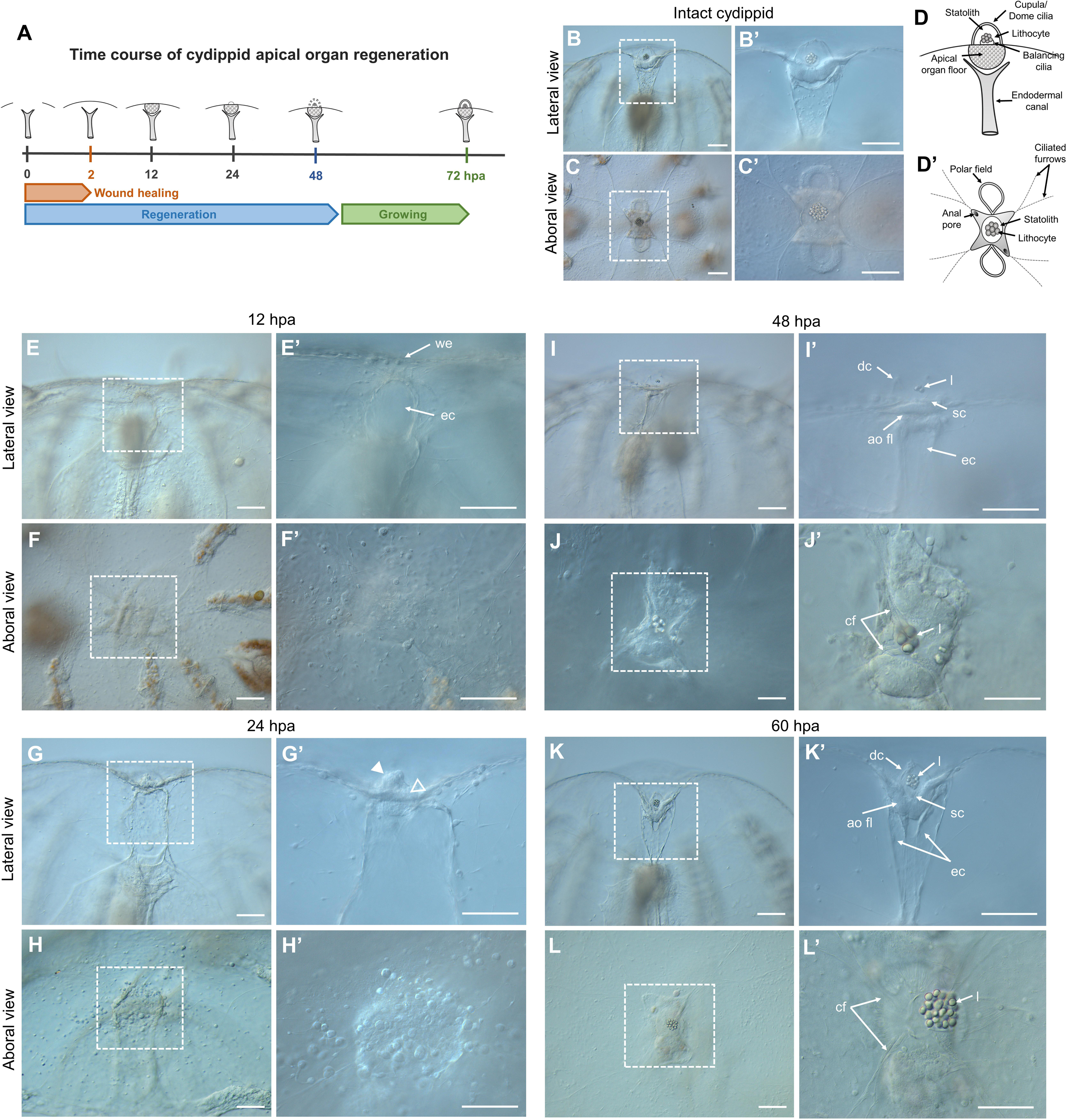
Apical organ regeneration in *Mnemiopsis leidyi* cydippids. (**A**) Schematic representation of the time course of morphogenic events during cydippid’s apical organ regeneration. Cartoons correspond to lateral views of the apical sensory organ at different stages of regeneration. (**B-C’**) DIC images of the apical organ of an intact cydippid in lateral (**B-B’**) and aboral (**C-C’**) views. (**D-D**) Schematic of the components of the apical sensory organ in lateral (**D**) and aboral (**D’**) views. (**E-L’**) DIC images showing the cut site of regenerating cydippids after apical organ amputation from 12 to 60 hpa. Lateral and aboral views are included. Dotted line rectangles delimit the area corresponding to higher magnifications on the right. Filled and empty white arrow caps in (**G’**) point to aggregation of cells forming the primordia of the future statolith and apical organ floor respectively. N>100. Scale bars = 100 μm. Abbreviations: hours post amputation (hpa), wound epithelium (we), endodermal canal (ec), dome cilia (dc), lithocyte (l), supporting cilia (sc), apical organ floor (ao fl), ciliated furrows (cf).

### Cell proliferation in intact cydippids

To identify areas of cell proliferation in juvenile *M. leidyi*, intact cydippids between 1.5 – 3.0 mm in diameter were labelled with the thymidine analog 5-ethynyl-20-deoxyuridine (EdU), which is incorporated into genomic DNA during the S-phase of the cell cycle (27,34–36) (Figure 5A). Cydippids incubated with EdU during a 15-minute pulse showed a pattern of cell division characterized by two main regions of active cell proliferation corresponding to the two tentacle bulbs (Figure 5B-B’). Higher magnifications of these structures showed EdU staining specifically concentrated at the lateral and median ridges of the tentacle bulb. Two symmetrical populations of densely packed cells were observed at the aboral extremity of the lateral ridges, previously characterized by Alié et al. (2011) as the aboral/external cell masses (a.e.c.) (Figure 5C’). EdU labeling was also found in some cells of the apical organ and few isolated cells along the pharynx and under the comb rows (n=20, Figure 5B-D’). To detect dividing cells in M phase of the cell cycle we performed anti-phospho-histone 3 (anti-PH3) immunolabelings in intact cydippids. The spatial pattern and distribution of PH3 labeling closely matched the one described for EdU incorporation, although PH3+ cells were always about 10% less numerous than the EdU labeled cells, suggesting that the duration of the M phase is much shorter than the S phase (n=10, Figure 5B’’,5C’’,5D’’).

**Figure 5.**
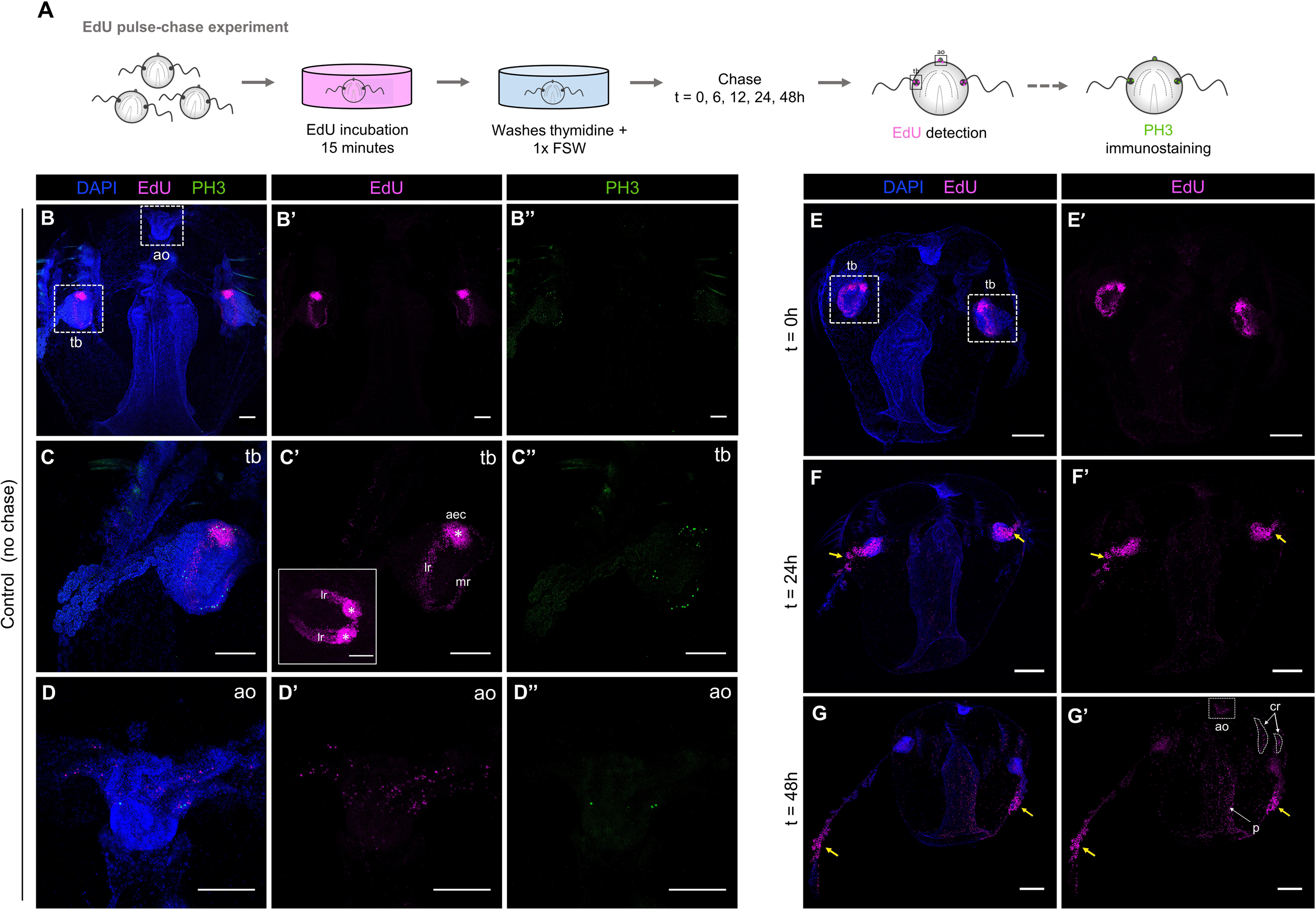
Cell proliferation in intact cydippids. (**A**) Schematic of the EdU pulse-chase experiment and PH3 immunostaining in intact cydippids. (**B-D’**) Confocal stack projections of whole intact cydippids oriented in a lateral view. White dotted rectangles in (**B**) delimit the tentacle bulb (tb) (**C-C’’’**) and apical organ (ao) (**D-D’’’**) structures showed in higher magnification at the bottom. Nuclei of S-phase cells are labeled with EdU (magenta), M-phase cells are immunostained with anti-phospho-Histone 3 (PH3) (green) and all nuclei are counterstained with DAPI (blue). Note that both markers of cell proliferation (EdU and PH3) show the same pattern of distribution along the cydippid’s body. The inset in (**C’**) shows an aboral view of the tentacle bulb after EdU staining. White asterisks in (**C’**) point to the symmetrical populations of intense cell proliferation referred as aboral/external cell masses (aec). (**E-G’**) Confocal stack projections of whole intact cydippids oriented in a lateral view. The time of the chase is listed at the top of the columns, and the labeling corresponding to each panel is listed to the left of the rows. Nuclei of S-phase cells are labeled with EdU (magenta) and all nuclei are counterstained with DAPI (blue). Note that EdU+ cells migrate from the tentacle bulb to the most distal end of the tentacle (yellow arrows). See **Additional file 2** for further detail of EdU pulse-chase experiment in the tentacle bulb. Scale bars = 100 μm. Abbreviations: apical organ (ao), tentacle bulb (tb), aboral/external cell masses (aec), lateral ridge (lr), medial ridge (mr), comb row (cr).

In order to track the populations of proliferating cells over time in intact animals we performed EdU pulse-chase experiments consisting in a 15-minute EdU incubation (pulse) and a chase of different times followed by visualization (Figure 5A). After a 24h chase, the pools of proliferating cells had migrated from the tentacle bulb through the proximal region of the tentacles, although some EdU+ cells were still detected at the tentacle sheath. Increased labeling of nuclei in the apical organ, pharynx and comb rows was also observed (n=10, Figure 5E-F’). Following a 48h chase, the population of proliferating cells that was originally in the tentacle bulbs at the time of labeling had migrated to the most distal end of the tentacles, but only a few cells associated with the tentacle bulb showed long-term EdU retention, suggesting that there is a resident population of slowly dividing stem cells in the tentacle bulb as previously reported by Alié et al. (2011). The number of EdU+ nuclei along the pharynx, the apical sensory organ (specifically in the apical organ floor) and comb rows was considerably increased compared to the 24h chase condition (n=10, Figure 5G-G’), suggesting that there are either small populations of EdU labeled cells restricted to those areas that had proliferated during the chase period, or that cells migrated in to those regions from regions of high mitotic density, or a combination of both events. Attempts to quantify and compare brightness levels of EdU positive cells to infer additional rounds of cell division during the chase period proved inconclusive (data not shown).

### Cell proliferation is activated during ctenophore regeneration

In order to determine the role of cell proliferation in ctenophore regeneration we performed a series of EdU experiments in regenerating cydippids. A 15-minute exposure to EdU at different times after surgical cutting was evaluated after two types of surgeries that required different regenerative responses: a bisection through the oral-aboral axis and an apical organ ablation (Figures 6A and 7A). The dynamics of cell proliferation at the wound site were quantified by calculating the ratio of EdU+ nuclei to total nuclei (DAPI stained) at different time-points following surgery (Figures 6B and 7B).

**Figure 6.**
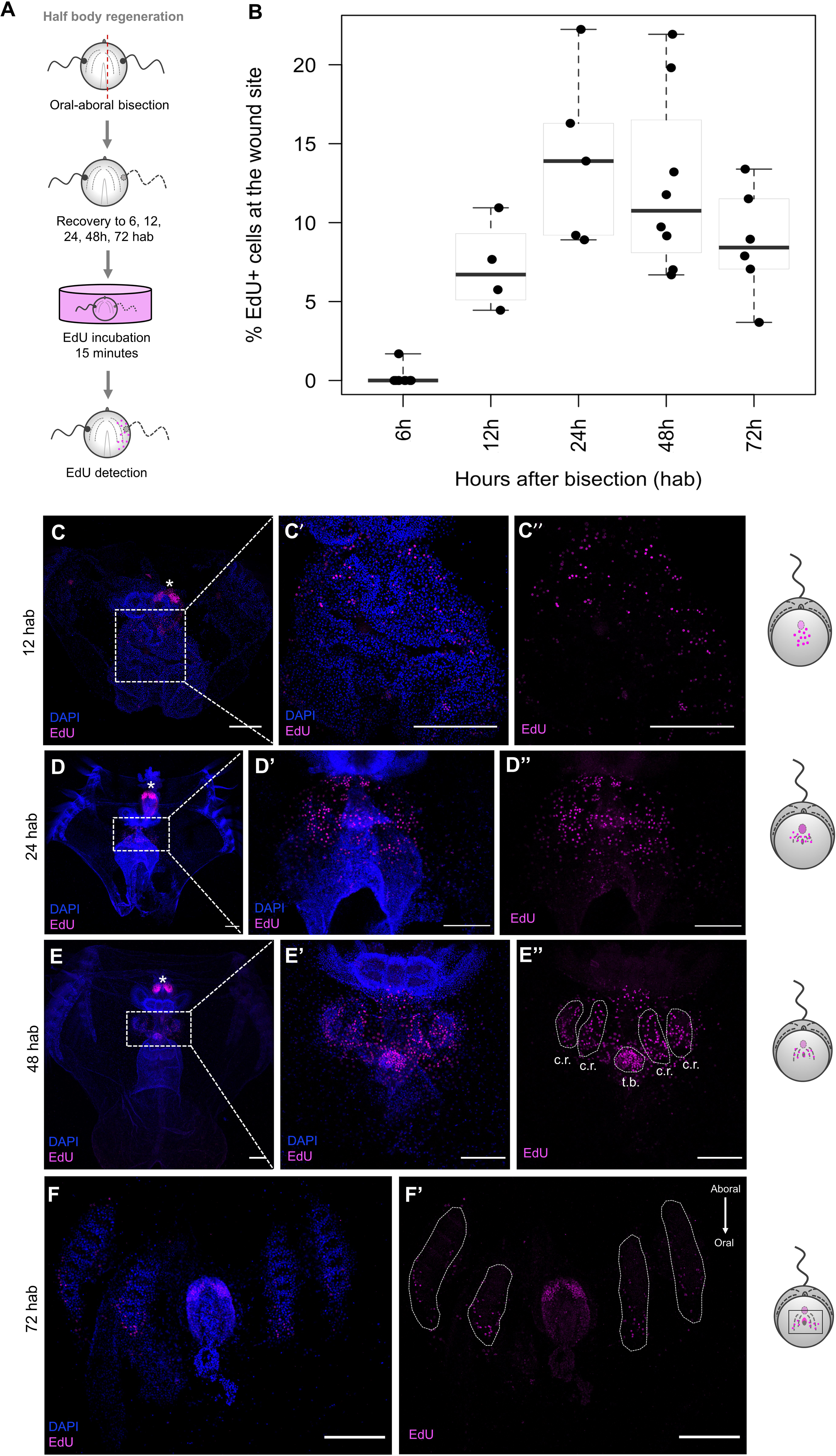
Cell proliferation during half body regeneration. (**A**) Schematic of the EdU incorporation experiment in cydippids bisected through the oral-aboral axis. (**B**) Box plot showing the levels of cell proliferation at the wound site at different time points after bisection. The thick horizontal bars indicate the median values. Each dot represents one individual. (**C-F’**) Confocal stack projections of bisected cydippids through the oral-aboral axis oriented in a lateral view showing the cut site in the first plane. The time following bisection is listed to the left of the rows. Nuclei of S-phase cells are labeled with EdU (magenta) and all nuclei are counterstained with DAPI (blue). The pattern of EdU labeling corresponding to each time-point is shown in a cartoon on the right of the rows. Dotted line rectangles in (**C**), (**D**) and (**E**) show the area corresponding to higher magnifications on the right. White dotted lines in (**E’’**) and (**F’**) delimit the area corresponding to the regenerating comb rows and tentacle bulb. White asterisks point to tentacle bulbs of the uncut site. Note that EdU+ cells at 72 hpa (**F’**) are located at the oral end of the regenerating comb rows and no EdU+ cells are detected at the aboral end where cells are already differentiated. Scale bars = 100 μm. Abbreviations: hours after bisection (hab) comb row (cr), tentacle bulb (tb).

**Figure 7.**
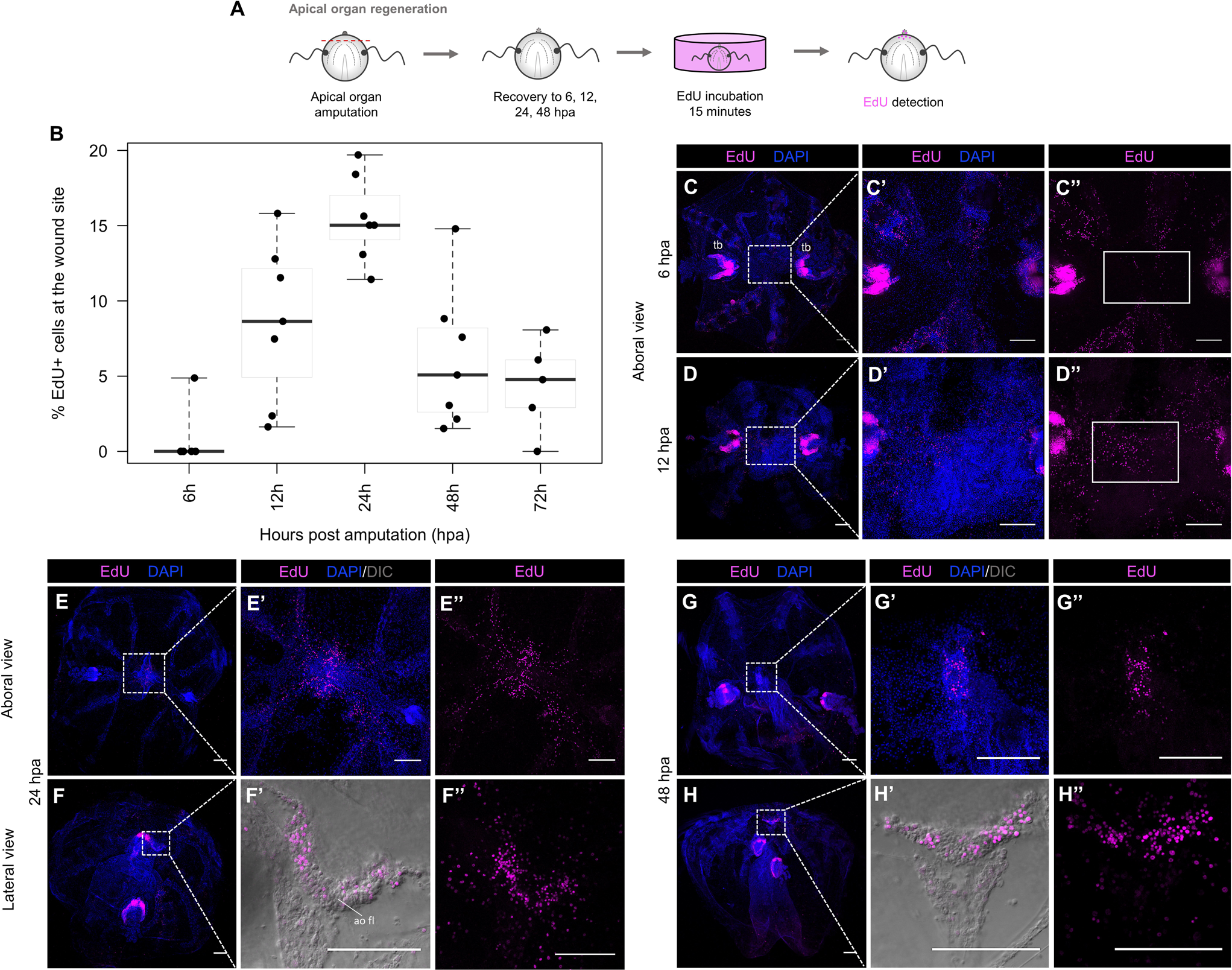
Cell proliferation during apical organ regeneration. (**A**) Schematic of the EdU incorporation experiment in cydippids in which the apical organ was amputated. (**B**) Box plot showing the levels of cell proliferation at the wound site at different time points post amputation. The thick horizontal bars indicate the median values. Each dot represents one individual. (**C-H’’**) Confocal stack projections of cydippids in which the apical organ was amputated at different time points post amputation. Aboral and lateral views are shown. The labeling corresponding to each panel is listed at the top of the columns, and the time following amputation is listed to the left of the rows. Dotted line rectangles in (**C**), (**D**), (**E**), (**F**), (**G**) and (**H**) show the area corresponding to higher magnifications on the right. White rectangles in (**C’**’) and (**D’’**) delimit the area of apical organ regeneration. Nuclei of S-phase cells are labeled with EdU (magenta) and all nuclei are counterstained with DAPI (blue). DIC images of the tissue are shown in (**F’**) and (**H’**). Scale bars = 100 μm. Abbreviations: tentacle bulb (tb), apical organ floor (ao fl), hours post amputation (hpa).

Following oral-aboral bisection, EdU+ nuclei were first detected at the wound site between 6 and 12 hours after bisection (hab). There was some variability in the presence of EdU+ nuclei at 6 hab – with some specimens having fewer EdU+ nuclei at the wound site than others – however the presence of EdU+ cells was consistent in all the analyzed individuals by 12 hab. The few EdU+ cells at the early stages were scattered all along the cut site, but no aggregation of cells was observed (n=7, Figure 6C-C’’). The number of EdU+ nuclei at the wound site slightly increased between 12 and 24 hab reaching a maximum at 24hab (Figure 6B), when EdU+ cells appeared concentrated in discrete areas corresponding to the forming primordia of the regenerating tissues (the tentacle bulb and comb rows) (n=27, Figure 6D-D’’). By 48 hab, the % of EdU+ nuclei had decreased as the cells started to differentiate into the final structures. EdU+ nuclei appeared confined into the regenerating comb rows and tentacle bulb, already distinguishable by nuclear staining (n=12, Figure 6E-E’’). At 72 hab, the number of EdU+ nuclei at the comb rows was considerably reduced and these were concentrated at the oral end of the regenerating structures, where oral portions of structures are generated later than aboral regions. For example, proliferative cells were no longer detected at the aboral end of the comb rows where cells had already differentiated into comb plates. In contrast, EdU+ cells at the regenerating tentacle bulb were abundant but appeared organized at the aboral extremity forming the two symmetrical populations of cells characteristic of the structure of the tentacle bulb (n=15, Figure 6F-F’). By 96 hab, when major repatterning events of regeneration were completed, EdU+ cells were only detected at the regenerated tentacle sheath forming the pattern of cell proliferation previously described in the tentacle bulbs of intact cydippids (Figure 5) (n=5, **Additional file 3A-A’’**). In combination with EdU incorporation experiments, anti-PH3 immunostaining was performed at selected time-points following bisection. PH3+ cells were detected in the regenerating comb rows and tentacle bulb at 24 hab and 48 hab (**Additional file 4A-B’’**) consistent with the EdU incorporation pattern, although the number of PH3+ cells was always less numerous than the EdU+ cells.

EdU labeling was also detected at the wound site of regenerating cydippids after apical organ amputation. Consistent with the oral-aboral bisection surgeries, EdU+ cells were first detected at 12 hpa suggesting that the start of the cell proliferation response occurred between the 6 and 12 hpa time points. A peak of cell proliferation was also observed at 24 hpa (Figure 7B), with EdU+ cells localized at the primordia of the apical organ, specifically in the apical organ floor and in the surrounding tissue including the regenerating comb rows adjacent to the cut site (n=15, Figure 7E-F’’). The number of proliferating cells slightly decreased at 48 hpa when EdU+ cells were concentrated in the regenerating apical organ and were no longer found in the tissues near the wound site (n=20, Figure 7G-H’’). By 72 hpa, the EdU+ nuclei were scarce and localized mostly along the polar fields in some specimens, while EdU+ nuclei were completely absent in other individuals at the same time-point (n=6, **Additional file 3B-C’’**). Anti-PH3 immunostaining showed presence of M-phase cells at the regenerating area at both 24 hpa and 48 hpa. Similar to half body regeneration, the pattern of anti-PH3 was consistent with the EdU labeling with PH3+ cells being more numerous at 24 hpa than 48 hpa. However, it should also be noted that as in all of our other observations, the number of PH3+ cells at any given time-point was always lower compared to the EdU+ cells (**Additional file 4C-D’**).

Interestingly, for both types of surgeries, proliferating cells were not organized in a compacted mass of “blastema-like” cells from were new tissue formed. In contrast, proliferating cells were very few and scattered throughout the wound site at early time-points after surgery – when a blastema is normally formed in animals with cell-proliferation based regeneration – and appeared more abundant and directly confined at the correct location of missing structures at later stages of regeneration, where they differentiated in place.

### Cells participating in the regenerative response appear to arise locally

To investigate the source of cells that contribute to the formation of new tissue during ctenophore regeneration we performed a series of EdU pulse and chase experiments in regenerating cydippids. This technique has been successfully used in different model systems as a strategy to indirectly track populations of proliferating cells and determine their contribution to the formation of new structures (19, 37). With the aim of determining whether cells proliferating before amputation contribute to the formation of new tissues, uncut cydippids were incubated in EdU, which was incorporated into cells undergoing the S-phase of cell cycle. After a 15-minute pulse, animals were removed from EdU, and EdU incorporation was blocked (“chased”) with several washes of thymidine and 1x FSW. Following the washes, apical organ amputations and oral-aboral bisections were performed and animals were left to regenerate in 1x FSW. The location of EdU+ cells was subsequently visualized at 24 and 48 hours after injury. In combination with EdU detection, an immunostaining against PH3 was performed in order to detect cells that were actively dividing in the animal immediately before fixation (Figure 8A).

**Figure 8.**
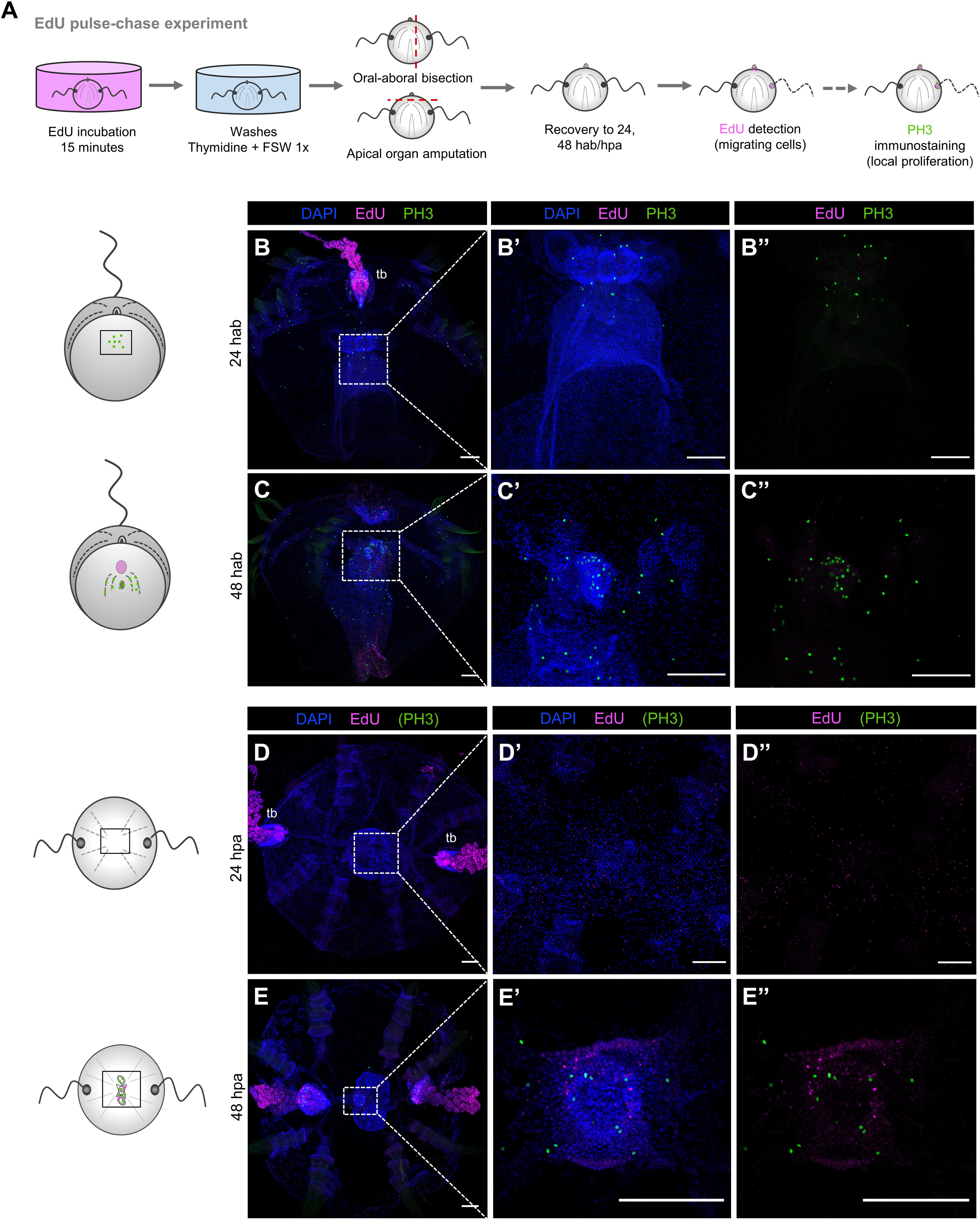
S-phase cells derived from the main regions of cell proliferation do not contribute to the formation of new structures. (**A**) Schematic of the EdU pulse-chase experiments and PH3 immunostaining in regenerating cydippids after oral-aboral bisection and apical organ amputation. (**B-C’’**) Confocal stack projections of bisected cydippids through the oral-aboral axis oriented in a lateral view showing the cut site in the first plane. (**D-E’’**) Confocal stack projections of cydippids in which the apical organ was amputated oriented in an aboral view. The labeling corresponding to each panel is listed at the top of the columns, and the time of chase is listed to the left of the rows. Nuclei of S-phase cells are labeled with EdU (magenta), M-phase cells are stained with anti-PH3 (green) and all nuclei are counterstained with DAPI (blue). Dotted line rectangles in (**B**), (**C**), (**D**) and (**E**) show the area corresponding to higher magnifications on the right. The pattern of EdU and PH3 staining is shown in cartoons on the left. Scale bars = 100 μm. Abbreviations: hours after bisection (hab), hours post amputation (hpa), tentacle bulb (tb).

No EdU+ cells were detected at the wound site at 24h (n=30) nor 48h (n=10) after bisection (Figure 8B-C’’). EdU labeling at the tentacle bulb resembling the pattern of cells migrating from the tentacle bulb distally along the tentacle previously described (Figure 5F-F’) confirmed that the chase worked properly (Figure 8B). Moreover, presence of PH3+ cells were observed at the regenerating area indicating active cell division at the moment of fixation (Figure 8B’’ and 8C’’). Following apical organ amputation, few EdU+ nuclei were detected at the area of apical organ regeneration although the EdU signal was very weak, suggesting that these cells were the result of multiple rounds of division (n=13, Figure 8D-D’’). After a 48h chase, few bright EdU+ nuclei were detected at the apical organ suggesting that S-phase cells from the uncut tissue might contribute to the formation of the apical sensory organ at later stages of regeneration (n=12, Figure 8E-E’’). Presence of PH3+ cells at the regenerating apical organ confirmed active cell division at the apical organ area (Figure 8E’-E’’). Taken together, these results show a minor contribution of proliferative cells originating in distant pre-existing proliferative tissue such as the tentacle bulbs to the formation of new structures.

Expression patterns determined through *in-situ* hybridization have reveled spatially restricted expression of the stem cell gene markers *Piwi, Vasa, Nanos and Sox* within areas of cell proliferation including the tentacle bulbs, in both juvenile cydippid and adult stages (27,28,36). On the other hand, the ctenophore group of Beroids do not possess tentacles at any stage of their life cycle and they are the only group of ctenophores that have lost the ability to regenerate (24). Based on these observations, it was hypothesized that the tentacle bulbs served as putative “stem cell niche” source of new cells involved in the regeneration process (23). To test this hypothesis, we surgically removed both tentacle bulbs from juvenile cydippids and assessed their ability to regenerate. Two days after amputation all animals had regenerated all the cell types of the tentacle bulb (n>100, Figure 9A-C’). EdU labeling at different time-points after amputation showed activation of cell proliferation during tentacle bulb regeneration, consistent with the other two types of surgeries analyzed. EdU+ nuclei were first detected at the distal end of the endodermal canals that connect the tentacles to the gastrovascular system at 18 hpa (n=10, Figure 9E-E’’). At 24 hpa the number of EdU+ cells had increased, and they were mainly organized forming the primordia of tentacle bulbs although some EdU+ cells were still detected at the tip of the endodermal canal connecting to the tentacle bulbs in formation (n=20, Figure 9F-F’’). By 48 hpa, EdU+ nuclei appeared organized in the characteristic pattern of intact tentacle bulbs (Figure 5B’ and 5C’), and they were not detected at the endodermal canals any more (n=20, Figure 9G-G’’). In addition, animals in which both tentacle bulbs and apical organ were removed, were able to regenerate all the missing structures (data not shown). These data argue strongly that the tentacle bulbs are not the source of multipotent stem cells required for the successful regenerative response in tentaculate ctenophores and point to a local source of cells in the formation of new structures.

**Figure 9.**
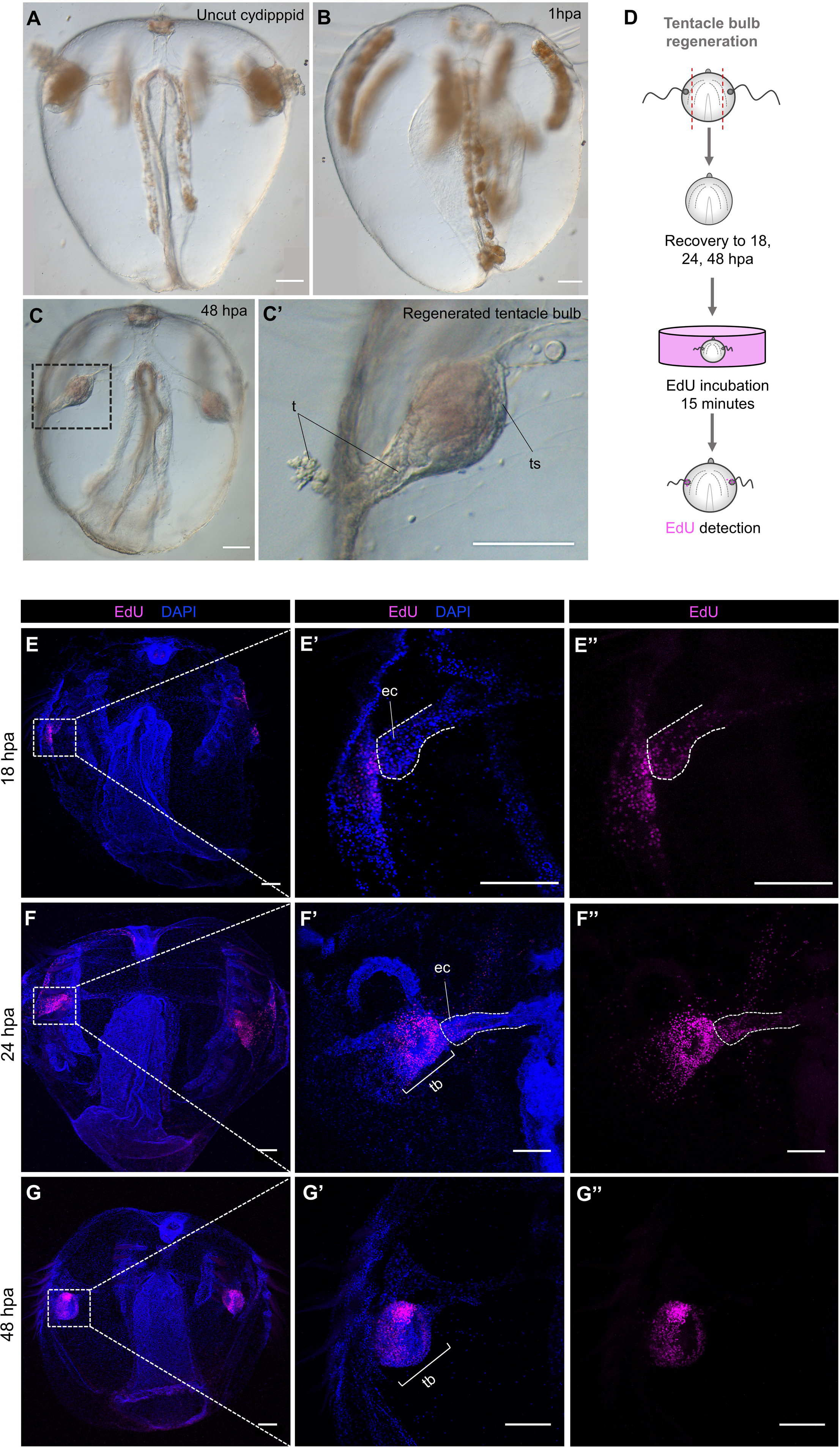
Regeneration occurs after removal of the main regions of active cell proliferation. (**A-C’**) DIC images of an uncut cydippid (**A**) and cydippids after tentacle bulbs amputation (**B-C’**). Black dotted line rectangle in (**C**) show the area corresponding to higher magnification on the right. (**D**) Schematic of the EdU incorporation experiment in cydippids in which both tentacle bulbs were amputated. (**E-G’’**) Confocal stack projections of cydippids in which the tentacle bulbs were amputated oriented in a lateral view at different time points post amputation. The labeling corresponding to each panel is listed at the top of the columns, and the time following amputation is listed to the left of the rows. Dotted line rectangles in (**E**), (**F**) and (**G**) show the area corresponding to higher magnifications on the right. Dotted lines in (**E’-E’’**) and (**F-F’’**) delimit the area corresponding to the endodermal canal. Nuclei of S-phase cells are labeled with EdU (magenta) and all nuclei are counterstained with DAPI (blue). Scale bars = 100μm. Abbreviations: tentacle sheath (ts), tentacle (t), endodermal canal (ec), tentacle bulb (tb), hours post amputation (hpa).

### There is recruitment of slowly-dividing cells at the regenerating structures

After having discarded the tentacle bulbs as the “stem cell niche” source of new cells for the regeneration of all missing structures, we questioned whether there is a population of slowly-dividing cells scattered among the ctenophore body which get stimulated after injury and contribute to the regenerative process. In order to target for slowly-cycling cells, we performed a long pulse-chase experiment consisting of a 2-hour EdU incubation pulse (in contrast to the 15-minute EdU pulse performed in previous pulse-chase experiments) followed by a 5-day chase (Figure 10A).

**Figure 10.**
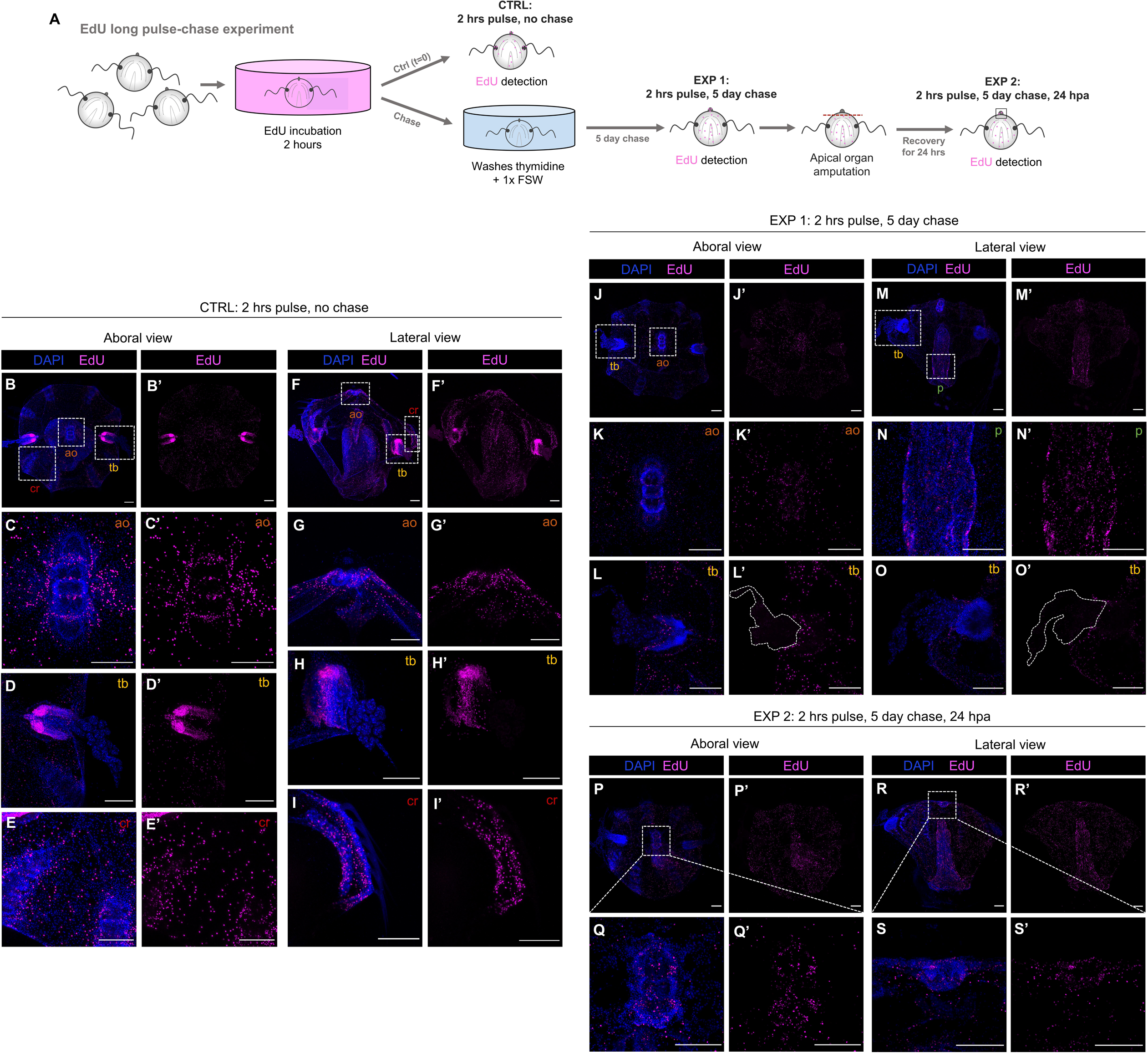
There is recruitment of slowly-dividing cells at the regenerating structures. (**A**) Schematic of the EdU long pulse-chase experiment. (**B-I’**) Confocal stack projections of whole intact cydippids oriented in an aboral (**B-E’**) and lateral (**F-I’**) views after 2-hour EdU pulse. White dotted rectangles in (**B**) and (**F**) delimit the apical organ (ao) (**C-C’** and **G-G’**), tentacle bulb (tb) (**D-D’** and **H-H’**) and comb row (cr) (**E-E’** and **I-I’**) structures showed in higher magnification at the bottom. (**J-O’**) Confocal stack projections of whole intact cydippids oriented in an aboral (**J-L’**) and lateral (**M-O’**) views after the 2-hour EdU pulse and 5-day chase. White dotted rectangles in (**J**) and (**M**) delimit the apical organ (ao) (**K-K’**), tentacle bulb (tb) (**L-L’** and **O-O’**) and pharynx (p) (**N-N’**) structures showed in higher magnification at the bottom. White dotted lines in (**L’**) and (**O’**) delimit the area corresponding to the tentacles. Note that no EdU+ cells are detected at the tentacle bulbs nor the tentacles after the 5-day chase (**L-L’**, **O-O’**) (see also **Additional files 10** and **11**). (**P-S’**) Confocal stack projections of cydippids in which the apical organ was amputated after the 2-hour EdU pulse and 5-day chase at 24 hpa. Aboral (**P-Q’**) and lateral (**R-S’**) views are shown. Dotted line rectangles in (**P**) and (**R**) show the area corresponding to higher magnifications on the bottom. Nuclei of S-phase cells are labeled with EdU (magenta) and all nuclei are counterstained with DAPI (blue). Scale bars = 100 μm. Abbreviations: hours post amputation (hpa), apical organ (ao), tentacle bulb (tb), comb row (cr), pharynx (p).

EdU+ cells just after the 2-hour EdU pulse were found very densely compacted in two main areas corresponding to the tentacle bulbs (Figures 10D-D’ and 10G-G’), and sparser although also abundant in the apical sensory organ (specifically the apical organ floor) (Figures 10C-C’, 10G-G’ and **Additional file 8**), under the comb rows (Figure 10I-I’) and the pharynx (Figure 10F’) (n=22, Figure 10B-I). This EdU labeling is consistent with the EdU pattern observed in intact cydippids after 15-minute EdU pulse (Figure 5B-B’), however, in contrast to the 15-minute pulse EdU pattern, the amount of EdU+ cells detected at the apical organ, comb rows and pharynx was considerably higher after the 2-hour EdU incubation. Moreover, EdU+ cells were also detected through the epidermal surface (**Additional file 6 and 7**) and endodermal canals (**Additional file 9**), locations where EdU+ cells were not detected after the 15-minute EdU pulse. After the 5-day chase, EdU+ cells were located around the apical organ, pharynx, comb rows and epidermis but they were no longer detected at the tentacle bulbs nor the tentacles (n=21, Figure 10J-O’ and **Additional file 10** and **11**). This result is consistent with the idea that a population of protected slowly-dividing cells does not exist confined in a concrete location (tentacle bulbs) but, rather, slowly-cycling cells are found scattered among several structures of the cydippid body.

To determine whether this population of slowly-dividing cells contribute to the process of regeneration, we amputated the apical organ structure from cydippids exposed to the 2-hour EdU pulse and 5-day chase. The location of EdU+ cells was subsequently visualized at 24 hpa (Figure 10A). EdU+ cells were detected at the regenerating apical organ 24 hours post amputation (n=10, Figure 10P-S’) indicating a contribution of slowly-dividing cells originated at the pre-existing tissue to the regenerating structure.

### Cell proliferation is strictly required for ctenophore regeneration

Having demonstrated that cell proliferation is activated during ctenophore regeneration, our next aim was to address the requirement of cell proliferation in the process of regeneration. Juvenile cydippids were exposed to hydroxyurea (HU) treatments, a drug that inhibits cell proliferation by blocking the ribonucleotide reductase enzyme and thereby preventing the S-phase of cell cycle (38). We first performed a dose-response test experiment in order to set the working concentration of HU in which animals could be continuously incubated during the complete period of regeneration with no significant disruption of their fitness. Concentrations of 20, 10 and 5mM HU were tested over a 72-hour time course. Incubations in 20 and 10mM HU were toxic and caused the degeneration and eventually death of most of the animals during the first 24 hours of incubation (data not shown). Incubations in 5mM HU were much less harmful; cydippids maintained a good condition, swimming normally with no cell death over the 72-hour time course. We therefore decided to set 5mM HU as the working concentration for the cell proliferation inhibitor experiments. We then assessed the efficacy of that drug concentration in blocking cell proliferation in intact cydippids. Intact cydippids were incubated in 5mM HU for 24 and 72 hours and then incubated for 15 minutes with EdU as previously described (**Additional file 12A**). At 24 hours of HU incubation, there was no detectable incorporation of EdU as compared with control cydippids, which showed the characteristic pattern of cell proliferation described in Figure 5 (**Additional file 12B-C’**). Inhibition of cell proliferation was maintained 72 hours after continuous HU incubation, as shown by the total absence of EdU+ cells in treated cydippids (**Additional file 12D-E’**). Finally, we evaluated the effect of the drug during regeneration in dissected cydippids. Cydippids bisected through the aboral-oral axis and cydippids in which apical organ was amputated were exposed to a continuous incubation of 5mM HU from 0 to 72 hours after surgery. None of the bisected cydippids had regenerated at 72 hours following HU treatment (n=75, Figure 11D-E’). Wound closure and healing occurred normally as shown by the continuous epidermal layer covering the wound (Figure 11E), but no sign of formation of the missing structures (tentacle bulb and comb rows) was observed. Moreover, bisected cydippids in which HU was added 4 hab (n=25) – when wound healing is already completed – and 12 hab (n=25) – when cells at the wound site have already begun to cycle – fail to regenerate the missing structures (data not shown). Likewise, none of the apical organ amputated cydippids had regenerated any of the structures/cell types of the missing apical organ at 72 hours following HU treatment, although the wound had correctly healed (n=55, Figure 11H-J’). Although HU treated animals failed to regenerate any of their missing structures, an aggregation of cells could be observed at the wound site (Figures 11E and 11J’). These accumulations of quite large round-shaped cells could correspond to undifferentiated cells ready to re-form the missing structures but not able to proceed due to the blocking of cell proliferation. Importantly, the absence of EdU incorporation in dissected cydippids treated with HU confirmed that cell proliferation was completely suppressed (**Additional file 12I-J’**). From these observations we conclude that regeneration was impaired due to the absence of cell proliferation, therefore, cell proliferation is indispensable for normal ctenophore regeneration.

**Figure 11.**
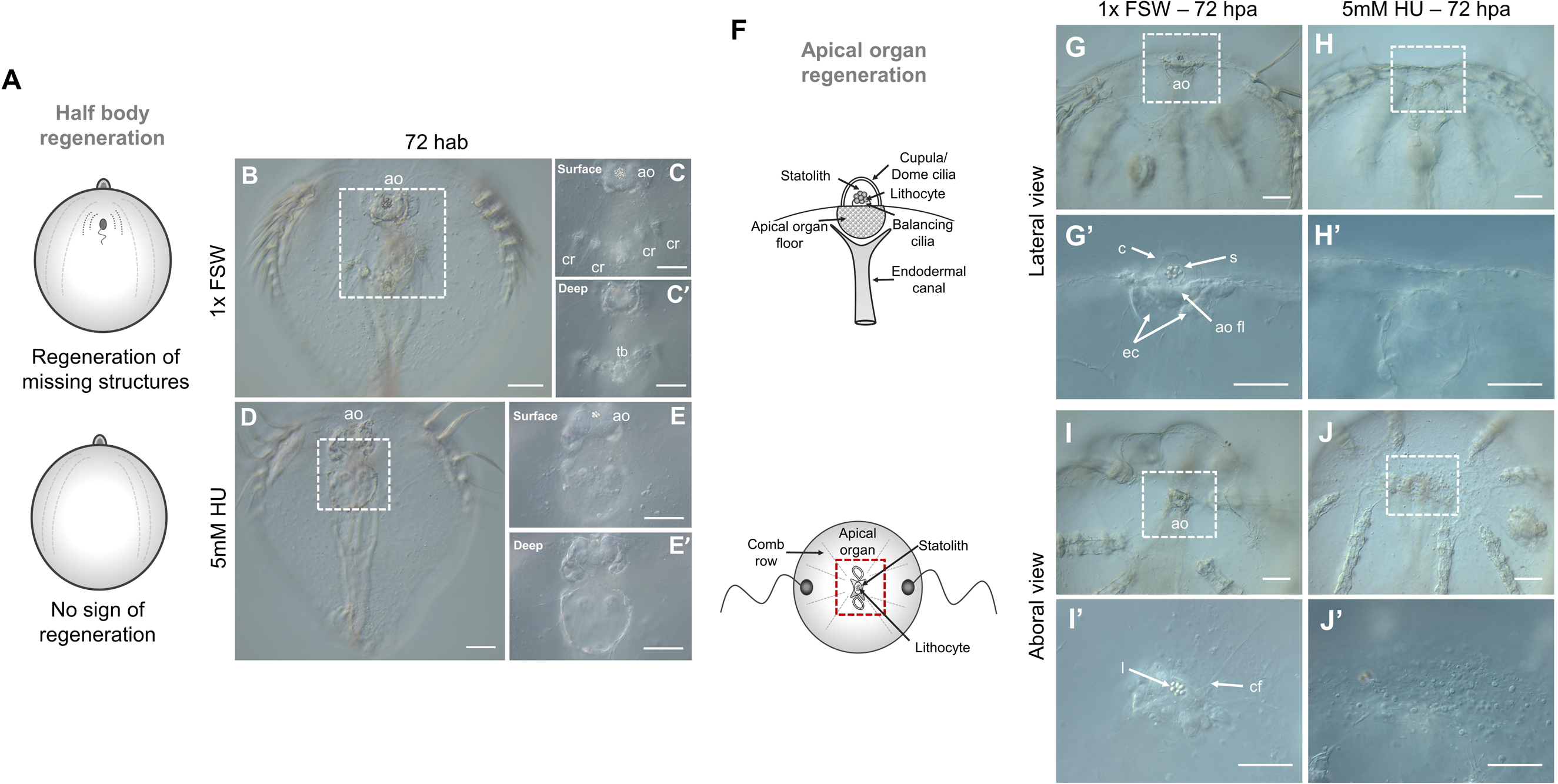
Ctenophore regeneration does not occur in the absence of cell proliferation. (**A**) Schematic representation of the regeneration state of cydippids shown in the panels on the right. Cartoons correspond to lateral views of the cydippid’s body bisected in half through the oral-aboral axis showing the cut site in the first plane. (**B-E’**) DIC images of bisected cydippids in a lateral view at 72 hab. The type of treatment corresponding to each panel is listed to the left of the rows. Dotted line rectangles in (**B**) and (**D**) show the area corresponding to higher magnifications on the right. Magnifications show surface (top) and deep (bottom) planes. Note that the wound site in treated cydippids is covered by a continuous epithelium but there is no sign of formation of missing structures. (**F**) Schematic representation of the apical sensory organ in lateral (top) and aboral (bottom) views. Red dotted rectangle at the bottom cartoon delimits the apical organ area shown in the images on the right. (**G-J’**) DIC images of cydippids in which the apical organ was amputated at 72 hpa orientated in a lateral (**G-H’**) and aboral (**I-J’**) views. The type of treatment corresponding to each panel is listed to the top of the columns. Dotted line rectangles in (**G**), (**H**), (**I**) and (**J**) show the area corresponding to higher magnifications on the bottom. Note that treated cydippids show aggregation of cells around the surface of the wounded area although none of the missing apical organ structures are formed. Scale bars = 100 μm. Abbreviations: hours after bisection (hab), hours post amputation (hpa), apical organ (ao), comb row (cr), tentacle bulb (tb), statolith (s), cupula (c), apical organ floor (ao fl), endodermal canal (ec), lithocyte (l), ciliated furrow (cf).

### Regenerative ability is recovered after HU treatment ends

Hydroxyurea has been shown to be reversible in cell culture following removal of the inhibitor (33) (Figure 12A). HU treatments on cut cydippids showed that wound healing occurs normally without cell division. In order to determine whether regeneration could be initiated in HU treated animals we took surgically cut cydippids that had been exposed to HU over 48 hours, washed them in 1x FSW to remove the inhibitor, and then followed their development for 48 hours to check for any ability to regenerate missing cell types (Figures 12B and 12E). Surprisingly, 36 out of 94 bisected cydippids (38%) had regenerated all the missing structures (comb rows, tentacle bulb and tentacle) 48 hours after HU had been removed (Figure 12D-D’’). 58 out of 94 bisected cydippids (62%) showed some signs of regeneration but ultimately remained as “half animals”, suggesting that these animals were not healthy enough to complete the regeneration process (40). (Note that these animals were not fed during the treatment (2 days) or recovery period (2 additional days)). On the other hand, 100% of the cydippids in which the apical organ was surgically removed and had been treated with HU for 48 hours, regenerated all the normal cell types of the apical organ (n=51, Figure 12H-I’). Altogether, these results show that ctenophore regeneration can be initiated over 48 hours after wound healing is complete, hence, wound healing and regeneration appear to be two relatively independent events which can be temporally decoupled.

**Figure 12.**
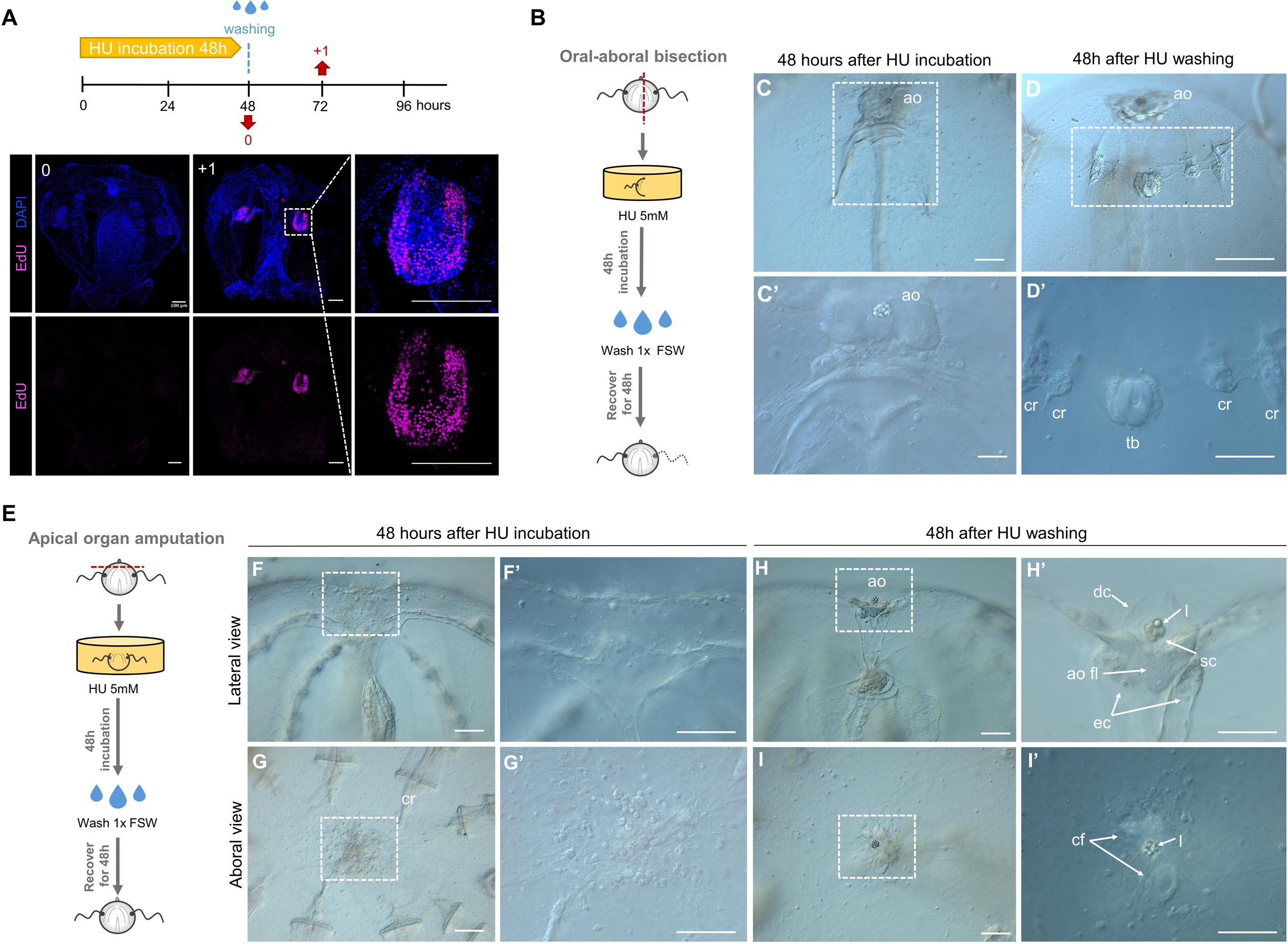
Regenerative ability is recovered after HU treatment ends. (**A**) Reversible S-phase arrest after HU treatment as detected by EdU labeling performed at the indicated time-points (red arrows). Nuclei of S-phase cells are labeled with EdU (magenta) and all nuclei are counterstained with DAPI (blue). Note that cells have already resumed cell cycle progression 1 day after HU wash. (**B** and **E**) Schematic representation of HU treatment and wash experiment in bisected cydippids through the oral-aboral axis (**B**) and cydippids in which the apical organ was amputated (**E**). (**C-C’**) DIC images of bisected cydippids in lateral view showing the wound site at 48 hab. Note that there is no sign of regeneration of the missing structures. (**D**-**D’’**) DIC images of bisected cydippids in lateral view showing the wound site at 96 hab. Note that all missing structures (comb rows and tentacle bulb) are regenerated. The dotted line rectangle in (**C**) and (**D**) show the area corresponding to higher magnification at the bottom. (**F-G’**) DIC images of amputated cydippids oriented in a lateral (top panels) and aboral (bottom panels) view showing the wound site at 48 hpa. Note that there is no sign of regeneration of the missing structures. (**H**-**I’**) DIC images of amputated cydippids oriented in a lateral (top panels) and aboral (bottom panels) view showing the wound site at 96 hpa. Note that all components of the apical organ are regenerated. Dotted line rectangles in (**F**), (**G**), (**H**) and (**I**) show the area corresponding to higher magnification on the right. Scale bars = 100 μm. Abbreviations: hours after bisection (hab), hours post amputation (hpa), apical organ (ao), comb row (cr), tentacle bulb (tb), dome cilia (dc), lithocyte (l), supporting cilia (sc), apical organ floor (ao fl), endodermal canal (ec), ciliated furrows (cf).

## DISCUSSION

In this study, we provide a detailed morphological and cellular characterization of wound healing and regeneration in the ctenophore *Mnemiopsis leidyi*. Wound closure is initiated immediately after injury, with the edges of the wound forming a round circumference that moves over the underlying mesoglea as it continues to reduce in diameter until they meet and forming a scar-less wound epithelium by 2 hours following injury. Two main mechanisms seem to be pivotal for ctenophore wound closure: active cell migration of cells from the mesoglea underneath the epithelium upwards to the edges of the wound; and dynamic extension of filopodia by the leading-edge epithelial cells in order to zipper the wound edges together. Cell migration and formation of actin-based cellular protrusions have been described during wound closure in multiple systems (41), however, slight differences in those mechanisms have been observed in ctenophore wound healing. First, cell migration takes place in a “deep to surface” direction instead of a lateral direction, suggesting that only specific cell-types from the mesoglea, such as mesenchymal cells, have the ability to migrate and contribute to gap closure. Second, wound-edge cells in ctenophores organize their cytoskeleton in spike-shaped filopodia rather than in plate-like extensions (lamellipodia), which happen to be the most common type of cellular protrusions among different model systems of wound healing, including the cnidarian Clytia (42). Despite these minor differences, the fact that common mechanisms of wound closure are shared between early branching phyla like ctenophores and cnidarians and bilaterians (including vertebrates) proves the ancient origin of wound healing mechanisms as a strategy to maintain epithelium integrity. Wound healing in *M. leidyi* takes place through changes in cell behavior and occurs normally in the absence of cell proliferation. This observation is consistent with the majority of animal models of regeneration found in cnidarians (12,20,42–44) as well as with the more phylogenetically distantly-related marine annelid worm *Platynereis dumerilii* (37). Following wound healing and prior to activation of cell proliferation in *M. leidyi*, there is remodeling of the tissue surrounding the wound and small numbers of round-shaped cells sparsely congregate at the wound site suggesting a reorganization of the tissue in order to prepare it for regeneration. Ctenophore regeneration, however, is strictly dependent on cell proliferation since none of the missing structures can be reformed in the absence of cell proliferation as proved by cell proliferation blocking treatments. Indeed, a combination of both tissue remodeling and cell proliferation-based strategies has been previously described in the regeneration of other animals including annelids (17, 45), although in those cases tissue remodeling takes place simultaneously with cell proliferation – or even subsequent to activation of cell proliferation – and is involved in the regeneration of a specific structures such as parapodia (46) or the gut (47).

Cell proliferation in *M. leidyi* is first detected at the wound site between 6-12 hours after surgery. The percentage of proliferating cells increases progressively during the first 12 hours following injury and reaches a maximum around 24 hours when the primordia of the missing structures are clearly delineated. Following this peak of cell proliferation, the percentage of cells undergoing cell division (S-phase) decreases while cells start to differentiate into their final structures. Comparing the kinetics of cell proliferation during regeneration of *M. leidyi* with the anthozoan cnidarian *Nematostella vectensis* (12), the percentage of dividing cells at the wound site is lower and the peak of maximum cell proliferation occurs earlier in ctenophore regeneration. In intact cydippids, cell proliferation is concentrated in two main areas of the cydippid’s body corresponding to the tentacle bulbs. Some actively cycling cells are also found in the apical organ as well as few isolated dividing cells along the pharynx and under the comb rows. These results are consistent with previous EdU analysis performed in *M. leidyi* cydippids (27, 28) and adult ctenophores of the species *Pleurobrachia pileus* (36) where EdU labeling has been detected in the same spatially restricted populations identified as stem cell pools, specialized in the production of particular cell types. Pulse-chase experiments show concrete areas of active cell proliferation in the tentacle bulb and progressive migration of these proliferating cells from the tentacle bulb to the distal tips of the tentacle. These observations fit with histological and cellular descriptions of the tentacle apparatus (36, 48) which identified different populations of undifferentiated progenitors source of all cell types found in the tentacle tissue. Surprisingly, long-term EdU retention is not detected in any of the cells of the tentacle apparatus suggesting that a population of “protected” slowly-dividing stem cells might not exist in the tentacle bulb. This opens the question of the nature and origin of the progenitor cells responsible for the maintenance of the homeostasis of the tentacle structure. Perhaps tentacle bulb stem cells are continuously recruited from adjacent somatic cells rather than being derived from a uniquely committed set of slow dividing “set aside” stem cells.

Interestingly, proliferating cells during regeneration do not organize to form a single large blastema-like structure from which a field of cells are reorganized to form the missing structures. Rather, small numbers of undifferentiated cells assume the correct location of all missing structures simultaneously and differentiate in place. Considering the early branching phylogenetic position of ctenophores in the tree of life (49, 50), the absence of blastema during ctenophore regeneration questions whether the formation of a blastema – which so far appears to have been reported in representatives of all phyla of regenerating animals (51) – is a conserved trait throughout the evolution of animal regeneration.

The strict requirement of cell proliferation and the absence of blastema formation could make ctenophore regeneration a case of non-blastemal cell proliferation-dependent regeneration. Although far less common than the blastemal based regeneration, isolated cases of non-blastemal regeneration have been reported such as lens regeneration by transdifferentiation in newts (52) or liver regeneration by compensatory proliferation in humans (53). EdU pulse-chase experiments after amputation show little to no contribution of cells originating in the main regions of active cell proliferation, including the tentacle bulbs, to the formation of missing structures. Moreover, the removal of these structures (tentacle bulbs), which have been reported to be localized areas of expression of genes involved in stem cell maintenance and regulation of cell fate (27,28,36) – and thus proposed to act as stem cell niches for regeneration – do not prevent regeneration. These observations argue against the contribution of discrete stem cell pools that migrate to and give rise to the re-formation of lost structures, suggesting that new structures are generated from a local source of cells that become activated to give rise to missing structures/cell types. Longer pulse-chase experiments in which the animals where incubated in EdU for an extended period of time (2 hours vs 15 min) and then followed by a much longer chase allowed the identification of a population of slowly-cycling (potentially stem) cells which could have escaped the initial short 15-minute pulse. Three main observations can be done from this experiment: 1) While the pattern of EdU+ proliferating cells after the long 2-hour pulse coincides with the one observed after the short 15-minute pulse, EdU+ cells after the chase are no longer found in any of the cells forming the tentacle apparatus. This observation is consistent with the results obtained in the pulse-chase experiments after amputation and thus strongly discards the existence of a stem cell niche in the tentacle bulbs source of cells for the regenerative process; 2) Long retaining EdU+ cells (referred as slow-dividing cells), are found uniformly distributed around the cydippid’s body instead of being organized in discrete pools. This distribution pattern of cell proliferation could be comparable to the neoblast distribution pattern characteristic of planarians (6,54,55), with the difference that neoblasts are known to reside in the planarian parenchyma – a mesenchymal tissue surrounding organ systems – (6), while these potential stem cells in ctenophores are found in both ectodermal and endodermal structures but not in the mesenchymal cells of the mesoglea. 3) EdU+ long retaining cells are detected at regenerating structures after amputation indicating that these slow-cycling cells contribute to the re-formation of new structures.

It is however important to note that our experiments do not give a definitive answer to the question of the origin of cells that give rise to new structures. There is the possibility that wound healing activates the dedifferentiation of cells at the wound site that are reprogrammed to give rise to whatever the appropriate set of cell types are needed to reconstitute the missing structures. The accumulation of large, round, apparently undifferentiated cells at the wound site during HU treatment is at least consistent with this scenario. On the other hand, wound healing could activate a dormant population of slowly-dividing pluripotent stem cells located uniformly around the body that could migrate to the wound site and drive the regeneration process, which could have escaped the short pulse of EdU incorporation and re-entered the cell cycle as a consequence of injury. Although we saw some evidence of cells migrating from the underlying mesoglea to repair the wound site, we did not see any evidence of long range cell migration to the site of new cell type formation. Nonetheless, an early study leveraging the combination of cell-lineage and specific cell-deletion experiments in *M. leidyi* showed that comb plate regeneration cannot occur when the entire complement of cell lineage comb plate progenitors are killed during embryogenesis, suggesting that, at least for comb plate regeneration, a semi-committed somatic stem cell population is set-aside during embryogenesis for comb plate replacement (32, 56). These data are premature and need to be extended to other cell types and later stages of the regenerative process, however the stereotyped cell lineage seen in ctenophores provides exciting opportunities to pursue the origins of stem cells in the regenerative process in living animals.

Overall, our data, together with evidences from previous studies in ctenophores, is consistent with a model of ctenophore regeneration based on a combined contribution of active cell proliferation at the wound site together with the recruitment of slowly-dividing cells to the regenerating structure (Figure 13). The fact that no discrete pools of neoblast-type stem cells are identified in the ctenophore body, and the absolute requirement for cell proliferation supports the hypothesis that proliferating cells at the wound site are the result of dedifferentiation events. On the other hand, whether these potential slowly dividing stem cells migrate to the wound site from close (local) or distant locations remains an open question. Gene expression data during the process of *M. leidyi* regeneration combined with cell tracing experiments will contribute to address these questions and thus refine our working model of the origin of cells during ctenophore regeneration. Molecular data during regeneration will also be very valuable for performing comparisons of gene expression profiles between *M. leidyi* development (57) and regeneration and thus determine whether the molecular basis of ctenophore regeneration is similar to that deployed during development.

**Figure 13.**
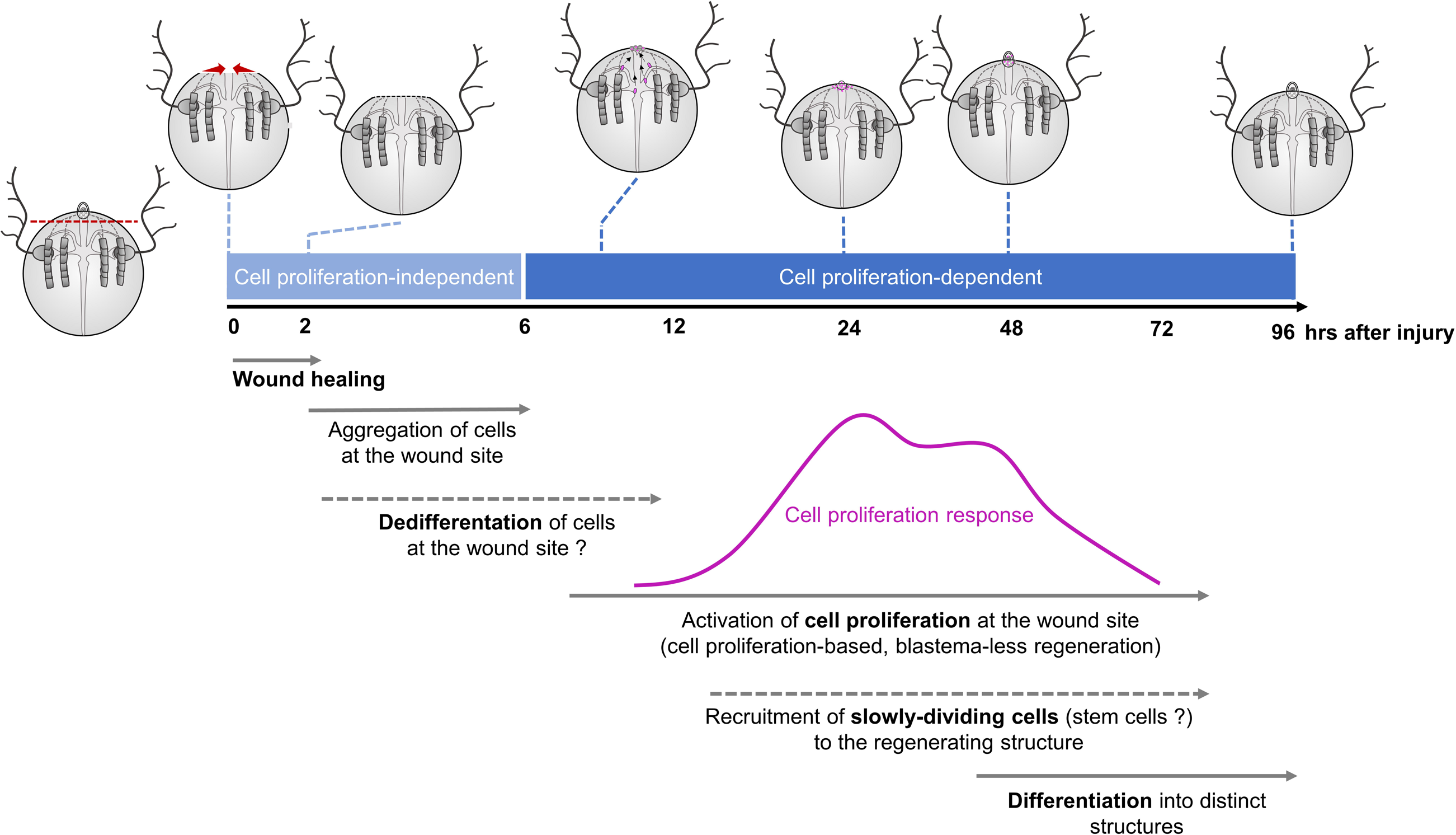
Working model of ctenophore regeneration. Timeline of the morphological and cellular events underlying ctenophore regeneration. Apical organ regeneration is used as example. Proliferating cells (EdU+) are colored in magenta. Wound closure is initiated immediately after surgery with the edges of the wound forming a round circumference that reduces in diameter until meeting and is completed within 1-2 hours after amputation. Reorganization of tissue including aggregation of round-shaped cells at the wound epithelium – potentially derived from dedifferentiation events – takes place during the first 6 hours after injury (Cell-proliferation independent phase). Cell proliferation is activated at the wound site between 6-12 hours after amputation and it reaches a maximum at 24 hours, when the primordia of the missing structures are formed. After this peak of cell proliferation, the number of proliferating cells at the wound site decreases while cells start to specify and differentiate into the final structures.

It is quite accepted that cells that re-epithelialize the wound (i.e the “wound epithelium”) provide the signals necessary to initiate regeneration (58, 59). In vertebrates, local thrombin activation is a signal for regeneration as shown by the study in which cultured newt myotubes returned to the cell cycle by the activity of a thrombin-generated ligand (60). On the other hand, cellular interactions also seem to be important for the initiation of the regenerative response. One such case is the dorsoventral interaction between the wounded tissues during wound healing in planarians which has been shown to play a key role in the formation of the blastema and, hence, initiation of regeneration (61). These observations suggest that wound healing and regeneration are two closely related processes which need to take place sequentially in time. Our results, however, show that ctenophore regeneration can be initiated over 48 hours after wound healing is completed, suggesting that regeneration can be initiated without direct signaling induced by the wounded epithelium. Regeneration of the missing structures is not initiated until the cell proliferation blocking treatment is removed. Hence, another case scenario is that the wound epithelium produces persistent signaling necessary for triggering regeneration at the time of wound healing, but the process cannot be initiated due to the blocking of cell proliferation. This is consistent with the proposed hypothesis for *Nematostella* that the key transition from wound healing to a state of regeneration is the activation of cell proliferation (62). Studying and comparing the molecular signaling involved in both ctenophore wound healing and regeneration will be very useful to get further insight into the relationship between these two processes.

## CONCLUSIONS

In conclusion, this study provides a thorough description of the morphological and cellular events during ctenophore wound healing and regeneration and compares them with the regenerative strategies followed by other metazoans. The early branching phylogenetic position of ctenophores together with their rapid, highly stereotyped development and remarkable ability to regenerate make them a key system to gain a better understanding of the evolution of animal regeneration.

## METHODS

### Animal care

Regeneration experiments were performed on juvenile *Mnemiopsis leidyi* cydippid stages due to their small size and ease of visualization and because their power of regeneration is the same as adults (30). *M. leidyi* cydippids were obtained from spawning adults collected from either the floating docks located around Flagler Beach area, FL. USA, or from the floating docks at the east end of the Bridge of Lions on Anastasia Island, St. Augustine, FL. USA. For spawning, freshly collected adults were kept in constant light for at least two consecutive nights and then individual animals transferred into 6” diameter glass culture dishes filled with 1x FSW and placed in total darkness. After approximately 3-4 hours in the dark at 22-24°C, these self-fertile hermaphroditic animals had spawned and embryos were collected by pipetting them into a new dish of UV treated 1.0 micron filtered full strength seawater (1x FSW) using a transfer pipette. Embryos were raised at 22-24°C for approximately 5-7 days and fed once a day with rotifers (*Brachionus plicatilis*, 160µm) (Reed Mariculture, Campbell, CA. USA).

### Animal surgeries

Operations were done in 35 mm plastic petri dishes with 2 mm thick silicon-coated bottoms (SYLGARD–184, Dow Corning, Inc.) on cydippids 1.5-3.0 mm in diameter. Cydippids were transferred in to the operation dishes in 0.2 µm-filtered seawater and cut using hand pulled glass needles from Pyrex capillaries (30). Three types of operations were performed: 1) Oral-aboral bisections, in which animals were cut longitudinally through the esophageal plane generating two “half-animals”. The operations were performed such that one half retained an intact apical organ while the remaining half lacked the apical organ. Only the halves retaining the apical organ were studied here as these halves regenerate to normal animals in a high percentage of the cases (30). 2) Apical organ amputations, involving the removal of the apical organ by cutting perpendicular to the oral-aboral axis above the level of the tentacle bulbs. 3) Tentacle bulb amputations, consisting in the removal of both tentacle bulbs (Figure 1D). Following surgery, halves containing the apical organ, amputated cydippids without apical organ and amputated cydippids without tentacle bulbs were returned to 35 mm plastic Petri dishes filled with 0.2 µm filtered 1x FSW for the desired length of time without feeding. All the regenerating experiments were performed at 22-24°C.

To study the wound healing process, juvenile cydippids were punctured generating a round-shaped wound of approximately 200-400 µm of diameter. Animals were placed in a small drop of water on a siliconized (Rain-X, Inc.) treated microscope slide and punctures were performed by pinching the epithelium layer using a pair of sharp needles (World Precision Instruments, Sarasota, FL. USA, Cat#500341). After puncture, animals were checked for the presence of an epithelial gap with the edges of the wound forming a small circumference exposing the mesoglea, and then they were immediately mounted for live imaging (see below).

### Tissue labeling and cell counts

#### Detection of cell proliferation by incorporation of EdU

To label proliferating cells, cydippids were fixed and processed for fluorescent detection of incorporated EdU using the Click-iT EdU labeling kit (Invitrogen Thermo Fisher Scientific, Waltham, MA. USA, Cat #C10424), which incorporates EdU in cells that are undergoing the S phase of the cell cycle. Specifically, intact cydippids between 1.5-3.0 mm in diameter or bisected/amputated cydippids were incubated in EdU labeling solution (100 μM of EdU in 1x FSW) for 15 minutes. For pulse-chase experiments cydippids were incubated with 100 μM EdU in 1x FSW for 15 minutes, washed 3 times with 100 μM thymidine in 1x FSW, and maintained in increasing volumes of 1x FSW until fixation. Control or operated cydippids were embedded in 1.2% low melt agarose (25°C melting temperature, USB, Inc Cat #32830) in a 35 mm plastic petri dish (Fisher, Inc. Cat #08757100A) and fixed in ice-cold 100mM HEPES pH 6.9; 0.05M EGTA; 5mM MgSO4; 200mM NaCl; 1x PBS; 3.7% Formaldehyde; 0.2% Glutaraldehyde; 0.2% Triton X-100; and 1x FSW (0.2 μm filtered) for 1 hour at room temperature with gentle rocking (63). Animals were then washed several times in PBS-0.02% Triton X-100, then one time in PBS-0.2% Triton X-100 for 20 min, and again several times in PBS-0.02% Triton X-100. The EdU detection reaction was performed according to manufacturer instructions using the Alexa-567 reaction kit. Following detection, cydippids were washed three times in PBS-0.02% Triton X-100, and subsequently all nuclei were counterstained with DAPI (Invitrogen, Carlsbad, CA. USA, Cat. #D1306) at 1.43 µM in 1x PBS for 2 hours. Cydippids were mounted in TDE mounting media (97% TDE: 970µl 2,2’-thiodiethanol (Sigma-Aldrich, St. Louis, MO. USA); 30µl PBS) for visualization. To quantify the percentage of EdU labeled cells at the wound site, Zeiss 710 confocal z-stack projections of operated cydippids were generated using Fiji software (Image J) and individual cells were digitally counted using Imaris, Inc. software (Bitplane, Switzerland). Only the area and z-stacks surrounding the wound site were used for the analysis. EdU+ cells and nuclei were counted separately in 5 to 10 specimens for each time-point. The number of EdU-positive nuclei were divided by the total number of nuclei stained with DAPI generating a ratio corresponding to the % of EdU+ cells.

#### Immunofluorescence

Proliferating cells in M phase were detected using an antibody against phospho Histone 3 (PH3 – phospho S10). Control or operated cydippids were fixed as mentioned above. Fixed cydippids were washed several times in PBS-0.02% Triton X-100 (PBT 0.02%), then one time in PBS-0.2% Triton X-100 (PBT 0.2%) for 10 min, and again several times in PBT 0.02%. They were then blocked in 5% normal goat serum (NGS; diluted in PBT 0.2%) for 1 hour at room temperature with gentle rocking. After blocking, specimens were incubated in anti-phospho histone H3 antibody (ARG51679, Arigo Biolaboratories, Taiwan) diluted 1:150 in 5% NGS overnight at 4°C. The day after, specimens were washed at least five times with PBS-0.2% Triton X-100. Secondary antibody (Alexa Fluor 488 goat anti-rabbit IgG (A-11008, Invitrogen, Carlsbad, CA. USA) was diluted 1:250 in 5% NGS and incubated over night at 4°C with gentle rocking. After incubation, specimens were washed three times with PBT 0.02% and incubated with DAPI (0.1μg/μl in 1x PBS; Invitrogen, Carlsbad, CA. USA, Cat. #D1306) for 2 hours to allow nuclear visualization. Samples were then rinsed in 1x PBS and mounted in TDE mounting media (97%TDE: 970µl 2,2’-thiodiethanol (Sigma-Aldrich, St. Louis, MO. USA); 30µl PBS) for visualization.

### Cell proliferation inhibitor treatment with Hydroxyurea (HU)

Cell proliferation was blocked using the ribonucleotide reductase inhibitor hydroxyurea (HU) (Sigma-Aldrich, St. Louis, MO. USA). Incubations with hydroxyurea were performed at a concentration of 5 mM in 1x FSW. Operated cydippids were exposed to continuous incubations of 5mM HU for 48-72 hours. HU solution was exchanged with freshly diluted inhibitor every 12 hours. For washing experiments, the effect of HU was reversed by removal and replacement of the drug with 1x FSW.

### Imaging

#### Visualization of living animals

In order to immobilize live animals for scoring and time lapse observations we utilized a custom made optically transparent jammed microgel. A solution of azobisisobutyronitrile (Sigma Aldrich, Inc.) and N’-Methylene Bisacrylamide (Sigma Aldrich, Inc) in ethanol is prepared at a 99:1 AAM:MBA molar ratio (64, 65). The solution is sparged with nitrogen for 30 minutes, then placed into a preheated 60°C oil bath. After approximately 30 minutes, the solution becomes hazy and a white precipitate begins to form. The reaction mixture is heated for an additional 4 hours, the precipitate collected by vacuum filtration and rinsed with ethanol on the filter. The microparticles are triturated with 500 mL of ethanol overnight. The solids are again collected by vacuum filtration and dried on the filter for ∼10 minutes. The particles are dried completely in a 50°C vacuum oven to yield a loose white powder. The purified microgel powder is dispersed in 0.2 µm-filtered seawater at a concentration of 7.5% (w/w) and mixed at 3500 rpm in a centrifugal speed mixer (64, 66) in five-minute intervals until no aggregates are apparent. The microgel is then left to swell overnight, yielding mounting medium made from packed hydrogel microparticles. The 7.5% microgel was placed around the operated/punctured cydippids mounted in a hydrophobic-treated slide (Rain-Ex, Inc). For short live imaging, a cover slip with clay corners was placed over the specimens; for long time-lapse live imaging, the corners of the cover slip were sealed using Vaseline in order to maintain the humidity of the preparation.

#### Microscopes and image analysis

*In vivo* differential interference contrast (DIC) images were captured using a Zeiss Axio Imager M2 coupled with an AxioCam (HRc) digital camera. Fluorescent confocal imaging was performed using a Zeiss LSM 710 confocal microscope (Zeiss, Gottingen, Germany) using either a 10x/0.3 NA dry objective, a 20x/0.8 NA dry objective or a 40x/1.3 NA water immersion objective. For time-lapse imaging, DIC images were captured using a Zeiss Axio Imager M2 coupled with a Rolera EM-C2 camera (Surrey, BC. Canada). Stacks were taken every minute. Generation of Z-stack projections, time-lapse movies and image processing was performed using Fiji software (67).

## ABBREVIATIONS

1x FSW: 1x filtered sea water
hab: hours after bisection
hpa: hours post amputation
EdU: 5-ethynyl-20-deoxyuridine
PH3: phospho-histone 3
HU: hydroxyurea
DIC: differential interference contrast.

## DECLARATIONS

### Acknowledgements

We thank the owners and staff of Marker 8 Hotel and Marina in St. Augustine for allowing us access to their floating docks for animal collection, and all the members of our lab for assistance and discussions.

### Availability of data and materials

All data generated or analysed during this study are included in this published article [and its supplementary information files].

### Authors’ contributions

Julia Ramon Mateu, Conceptualization, Investigation, Writing—original draft, Writing— review and editing; Tori Ellison and Thomas E. Angelini, design and synthesis of microgels; Mark Q. Martindale, Conceptualization, Supervision, Writing—original draft, Writing—review and editing. All authors read and approved the final manuscript.

### Ethics approval and consent to participate

Not applicable.

## ADDITIONAL FILE LEGENDS

**Additional file 1: Movie 1. Wound healing time-lapse.** DIC microscopy time-lapse movie of wound closure after puncture of the wound pictured in Figure 2. Note the migration of mesogleal cells to the edges of the wound during the wound closure process. Scale bar = 100 μm. [AVI, 13.6MB]

**Additional file 2: Figure S5. EdU pulse and chase experiment in the tentacle bulb.** (**A-D’**) Confocal stack projections of tentacle bulbs and tentacles in lateral view. The time of the chase is listed to the left of the rows, and the labeling corresponding to each panel is listed at the top of the columns. Nuclei of S-phase cells are labeled with EdU (magenta) combined with a differential interference contrast image of the tissue (DIC). Scale bars = 50 μm. [TIFF, 8.2MB]

**Additional file 3: Figure S6 and S7. EdU staining at 96 hab and 72 hpa. (A-A’’)** Confocal stack projections of a bisected cydippid through the oral-aboral axis at 96 hab oriented in a lateral view showing the cut site in the first plane. White asterisks point to tentacle bulbs of the uncut site. (**B-C’’**) Confocal stack projections of cydippids in which the apical organ was amputated at 72 hpa. Examples of the two types of EdU pattern observed at this time-point are shown. Nuclei of S-phase cells are labeled with EdU (magenta) and all nuclei are counterstained with DAPI (blue). The pattern of EdU labeling corresponding to each time-point is shown in a cartoon on the left of the rows. Scale bars = 100 μm. Abbreviations: hours after bisection (hab), hours post amputation (hpa), polar field (pf). [TIFF, 5.1MB]

**Additional file 4: Figure S6 and S7. PH3 immunostaining after oral-aboral bisection and apical organ amputation. (A-B’’)** Confocal stack projections of a bisected cydippid through the oral-aboral axis oriented in a lateral view showing the cut site in the first plane. The time following surgery is listed to the left of the rows. M-phase cells are stained with anti-PH3 (green) and all nuclei are counterstained with DAPI (blue). (**C-D**’) Confocal stack projections of regenerating apical organs in a lateral view. The time following surgery is listed to the left of the rows. M-phase cells are stained with anti-PH3 (green) combined with DIC image of the tissue. DIC images of surface and deep planes are shown for 24 hpa. Scale bars = 100 μm. Abbreviations: hours after bisection (hab), hours post amputation (hpa). [TIFF, 3.1MB]

**Additional file 5: Figure S8. Controls of the EdU pulse-chase experiments in regenerating cydippids.** (**A**) Diagram of the experimental setup. (**B-D’**) Confocal stack projections of an uncut cydippid just after EdU incubation (control 1). (**E-F’**) Confocal stack projections of an amputated cydippid just after the cut (control 2). Nuclei of S-phase cells are labeled with EdU (magenta) and all nuclei are counterstained with DAPI (blue). White dotted rectangles in (**B**) and (**E**) show the area corresponding to higher magnifications on the right. White arrows in (**E’**) point to the wound site. Note that no EdU+ cells are detected at the wound site just after the cut while EdU staining is found in the normal areas of cell proliferation (tentacle bulbs). Scale bars = 100μm. [TIFF, 3.6MB]

**Additional file 6: Movie S10. Z-stack of whole body intact cydippid oriented in an aboral view after a 2-hour EdU incubation pulse.** Nuclei of S-phase cells are labeled with EdU (magenta) combined with a differential interference contrast image of the tissue (DIC). Note that EdU+ cells are located at the surface epidermis but not at the mesenchymal cells of the deep mesoglea. Scale bar = 100μm. [AVI, 36.4MB]

**Additional file 7: Movie S10. Z-stack of whole body intact cydippid oriented in a lateral view after a 2-hour EdU incubation pulse.** Nuclei of S-phase cells are labeled with EdU (magenta) combined with a differential interference contrast image of the tissue (DIC). Scale bar = 100μm. [AVI, 38.9MB]

**Additional file 8: Movie S10. Z-stack of the apical organ area of a cydippid oriented in an aboral view after a 2-hour incubation pulse.** Nuclei of S-phase cells are labeled with EdU (magenta) combined with a differential interference contrast image of the tissue (DIC). Note that most of the EdU+ cells are found at the epidermis surrounding the apical organ area and some EdU+ cells are located at the apical organ floor. Scale bar = 100μm. [AVI, 12.8MB]

**Additional file 9: Movie S10. Z-stack of the comb row area of a cydippid after a 2-hour incubation pulse.** Nuclei of S-phase cells are labeled with EdU (magenta) and all nuclei are counterstained with DAPI (blue), combined with a differential interference contrast image of the tissue (DIC). Note that EdU+ cells are found at the endodermal canals that connect the comb rows with the digestive system. Scale bar = 100μm. [AVI, 17,7MB]

**Additional file 10: Movie S10. Z-stack of the tentacle apparatus of a cydippid oriented in an aboral view exposed to a 2-hour EdU pulse followed by a 5-day chase.** Nuclei of S-phase cells are labeled with EdU (magenta) combined with a differential interference contrast image of the tissue (DIC). Note that no EdU+ cells are found at the tentacle bulb nor the tentacle. Scale bar = 100μm. [AVI, 5.4MB]

**Additional file 11: Movie S10. Z-stack of the tentacle apparatus of a cydippid oriented in a lateral view exposed to a 2-hour EdU pulse followed by a 5-day chase.** Nuclei of S-phase cells are labeled with EdU (magenta) combined with a differential interference contrast image of the tissue (DIC). Note that no EdU+ cells are found at the tentacle bulb nor the tentacle. Scale bar = 100μm. [AVI, 1.5MB]

**Additional file 12: Figure S11. 5mM Hydroxyurea (HU) blocks cell proliferation in intact and bisected cydippids.** (**A**) Schematic of cell proliferation inhibitor experiments with HU in intact cydippids. Confocal stack projections of intact cydippids oriented in an aboral view at 24 hours (**B-C’**) and 72 hours (**D-E’**) after HU incubation. The type of treatment is listed at the top of the columns, and the labeling corresponding to each panel is listed to the left of the rows. Nuclei of S-phase cells are labeled with EdU (magenta) and all nuclei are counterstained with DAPI (blue). Note that no EdU+ nuclei were detected in treated cydippids after HU incubation. (**F**) Schematic of cell proliferation inhibitor experiments with HU in dissected cydippids. (**G-H’**) Confocal stack projections of an untreated bisected cydippid oriented in a lateral view at 72 hab. (**I-I**’) Confocal stack projections of a bisected cydippids treated with HU at 72 hab. (**J-J’**) Confocal stack projections of cydippids amputated from the apical organ treated with HU at 72 hpa. The type of treatment and the time following bisection are listed at the top of the columns, and the labeling corresponding to each panel is listed to the left of the rows. The dotted line rectangle in (**G**) shows the area corresponding to higher magnifications on the right. The white asterisks in (**G**), (**G’**) and (**I**) point to the tentacle bulb of the uncut site. The white rectangle in (**J**) indicates the area of apical organ regeneration. Nuclei of S-phase cells are labeled with EdU (magenta) and all nuclei are counterstained with DAPI (blue). Note that no EdU+ nuclei were detected and none of the missing structures had regenerated in treated cydippids. Scale bars = 100 μm. Abbreviations: hours after bisection (hab), hours post amputation (hpa). [TIFF, 4.3MB]

