## Supplementary figures and images for "Regeneration in the absence of a blastema requires cell division but is not tied to wound healing in the ctenophore *Mnemiopsis leidyi*"

### Sup. Fig. 2

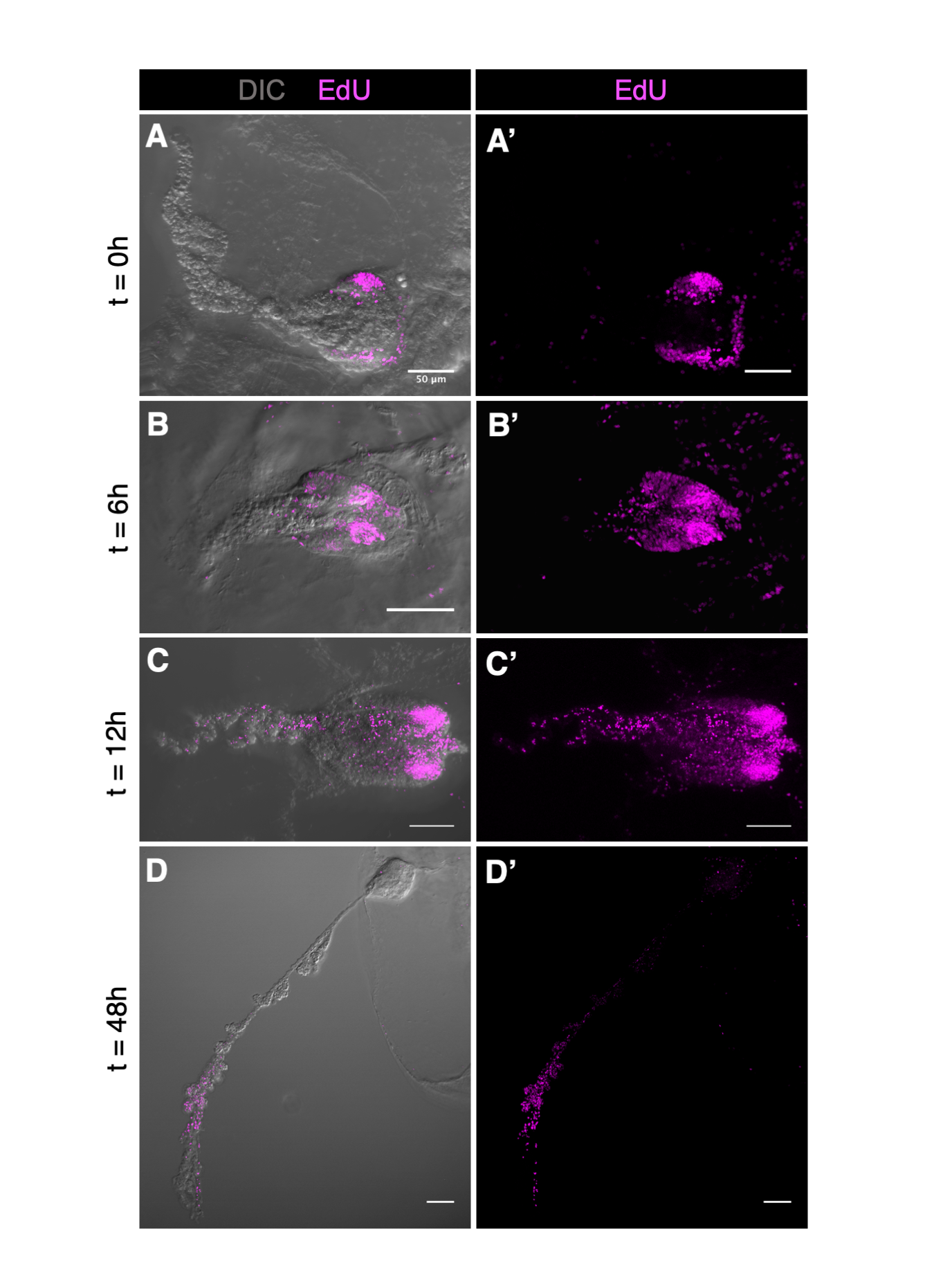

### Sup. Fig. 3

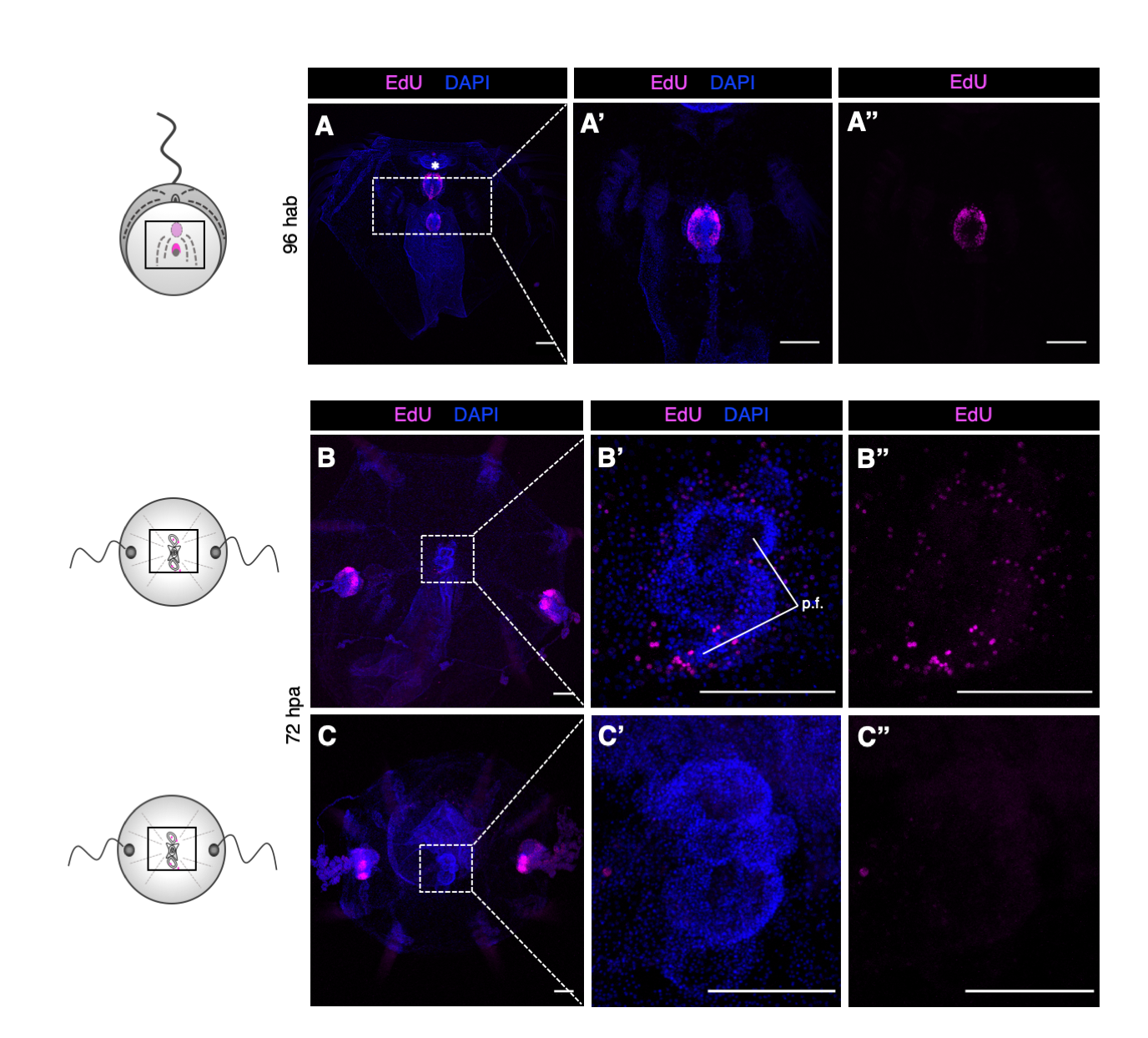

### Sup. Fig. 4

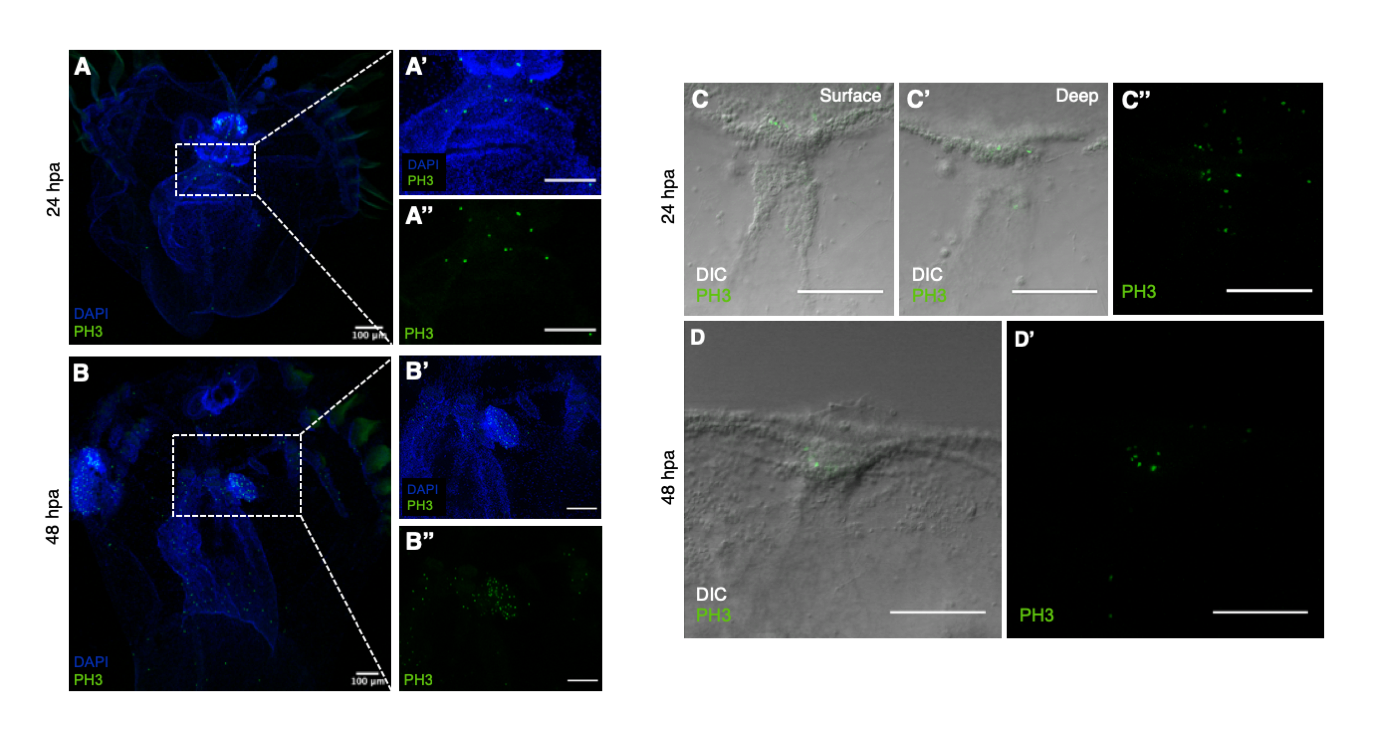

### Sup. Fig. 5

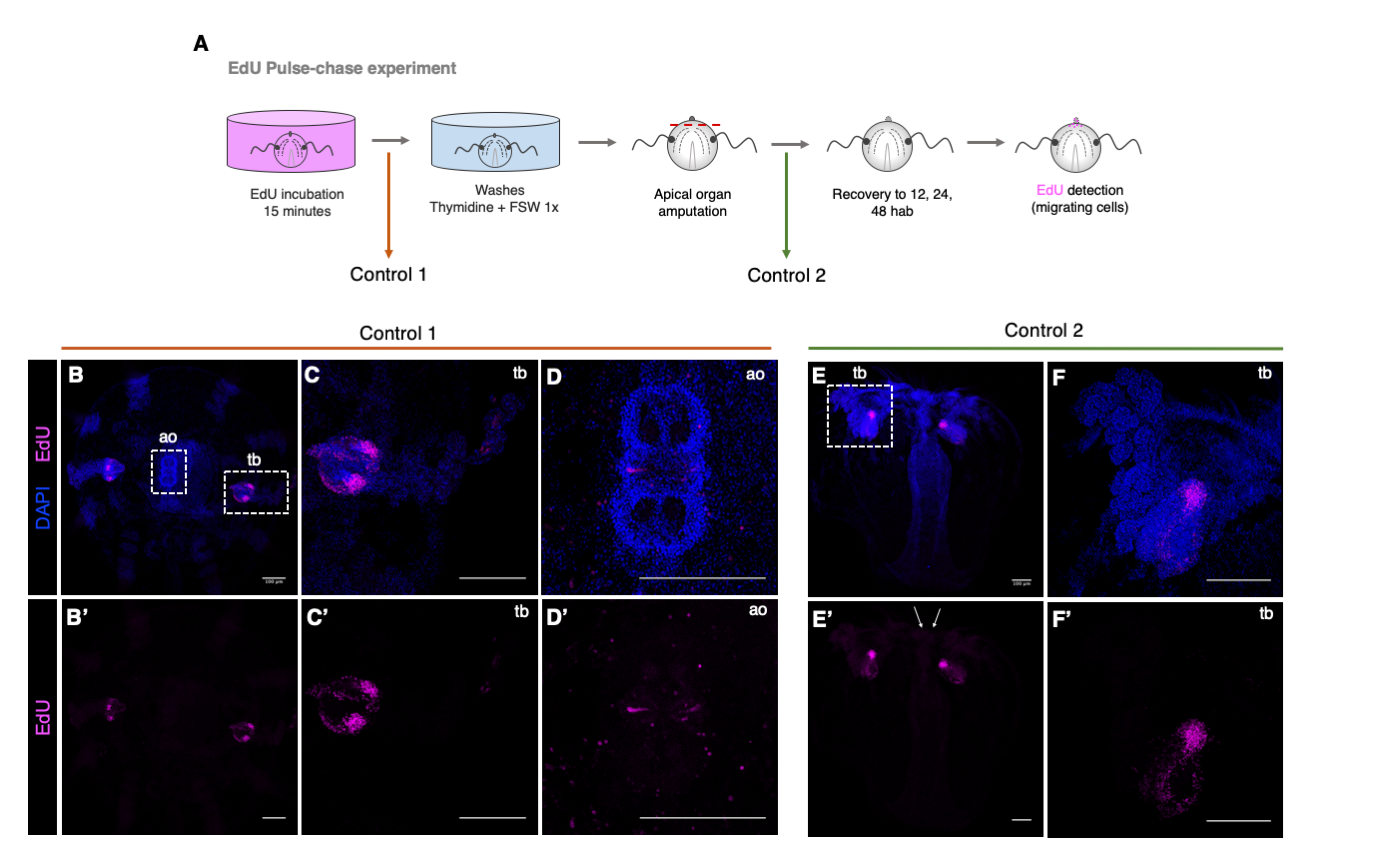

### Sup. Fig. 12

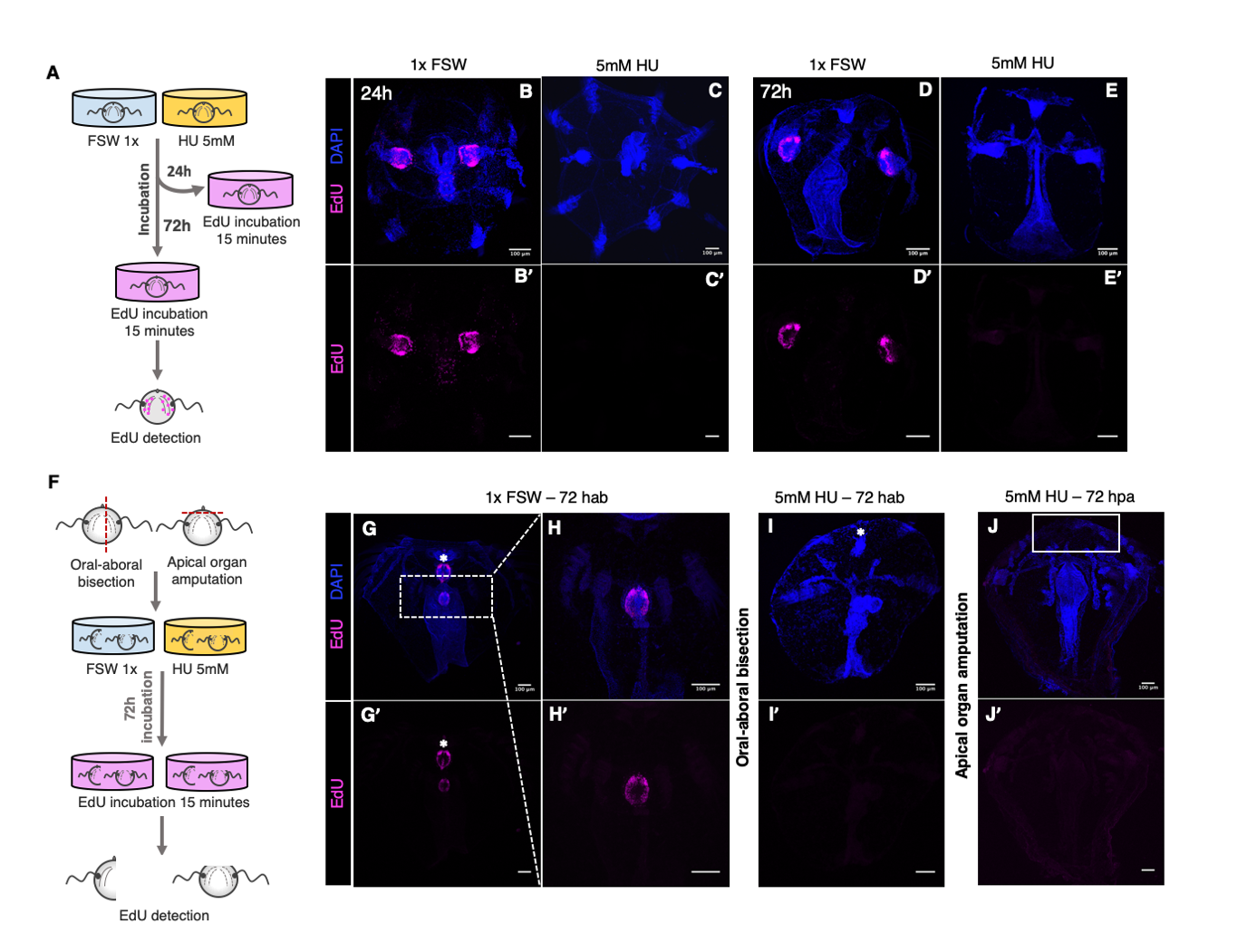
